# A Nature-Inspired Ion Trap for Parallel Manipulation of Ions on a Massive Scale

**DOI:** 10.1101/2025.08.21.671534

**Authors:** Andrew N. Krutchinsky, Brian T. Chait

## Abstract

Parallelization has revolutionized computing and DNA sequencing but remains largely unexploited in mass spectrometry (MS), which typically analyzes ions sequentially. We introduce a nature-inspired ion trap (MultiQ-IT) that enables massively parallel MS. The device comprises a cubic array of small quadrupoles forming multiple ion entry and exit ports, allowing >10⁹ ions to be confined and manipulated simultaneously. This architecture enables selective depletion of singly charged ions in real time, greatly improving signal-to-noise ratios and detection sensitivity. The trap also functions as a parallel ion splitter, transmitting ions into multiple m/z-specific beams. We demonstrate scalable ion throughput, real-time charge discrimination, and parallel beam separation, suggesting a path toward truly parallel MS. Our results offer a foundation for next-generation, high-throughput proteomic and metabolomic analyses.

## Main Text

Mass spectrometry (MS) is currently practiced predominantly as a sequential technique, wherein various species in a sample are selected and interrogated one after another (*1–3*). Because of the finite time needed to examine each species in turn, such sequential mode MS analysis suffers from inescapable limitations in sensitivity, speed and the ability to exhaustively analyze all ions produced from the sample, especially when the composition of the ion beam is complex, rapidly changing, highly variable, and of low abundance. The resulting loss of sensitivity makes comprehensive analysis of, e.g., the proteomes of single cells through peptide MS fingerprinting (*4, 5*) challenging with current approaches. One promising way to address these limitations is through the parallelization of MS analysis, which has the potential to greatly enhance throughput and sensitivity. This concept finds an analogue in parallel computing, where the Gustafson-Barsis law (*6*) has demonstrated the great power of parallelization; but the technology for efficient parallel MS analysis has remained underdeveloped, with a few notable exceptions (e.g., (*7, 8*)). In response to this challenge, we have designed and constructed a new type of ion trap (MultiQ-IT) intended for parallel manipulation of ions with the potential for application on a massive scale (*9–12*).

The MultiQ-IT was inspired by our research on nucleocytoplasmic transport through nuclear pore complexes (NPCs) and our proposed virtual gating mechanism (*13, 14*). We envisioned an ion trap analogue of the nucleus, where diffusion serves as the fundamental transport mechanism. However, rather than being embedded with NPCs, this system is engineered with an array of ion input and output ports, enabling ion confinement, controlled transport, and parallel processing. In this design, the ion confinement region is enclosed by a multitude of cylindrical electrodes, systematically arranged around a cube (**Fig. 1A,B; fig. S1**). These electrodes are driven by radio frequency (RF) signals (**fig. S2**), so that each group of four neighboring cylinders forms a quadrupole. This arrangement of RF quadrupoles, which we term a “MultiQ-Ion Trap (MultiQ-IT)”, serves to confine ions within the cubic structure by the pondermotive forces generated by the RF quadrupole arrays (*15, 16*). **Fig. 1C** shows an exemplar computed trajectory of a single ion injected into a MultiQ-IT that contains 486 quadrupoles (486Q). The trap operates at a pressure ∼0.1 Pa of nitrogen gas, so that after injection, ions are rapidly thermalized (usually within a few milliseconds) and begin diffusing inside the ion confinement region. Here, they periodically undergo bouts of re-excitation when ions diffuse into the proximity of the quadrupoles, followed by re-thermalization. At the elevated RF amplitude shown, the quadrupoles do not allow passage of the ions through their gaps because the Mathieu stability parameter chosen (*q=1.28)* lies outside of the *0<q<0.9* stability region (*17*). As we decrease the RF amplitude, ions can occupy a larger volume within the trap, and after thermalization can defuse between the quadrupoles, whereupon they can escape from the trap along the quadrupole axes (**Fig. 1D**). Their escape can be prevented either by elevating the potential difference between the quadrupoles and the exterior wall electrodes *(ΔU=U_wall_ - U_0_*) or encouraged by providing specific conditions for ion transport through the quadrupoles and the exterior walls. The conditions for ion transport imposed on each input/output quadrupole port can range from simple, as in a small static potential barrier that prevents thermalized ions from escaping the storage region, to more complex, which may involve operating each quadrupole as a separate filter capable of only transmitting ions with very specific m/z values (*18–20*). To explore the detailed behavior of MultiQ-ITs experimentally, we constructed several versions of the instrument (**Fig. 1E**) and, over the course of our research, replaced the commercial electrospray sources and interfaces of three different mass spectrometers with our own electrospray interface and MultiQ-IT systems (**Fig. 1E**). These instruments were used to investigate key properties of the MultiQ-IT, including ion capacity, ion residence time, and the feasibility of using multiple ion trap outputs for the parallel manipulation of ions.

**Fig. 1.**
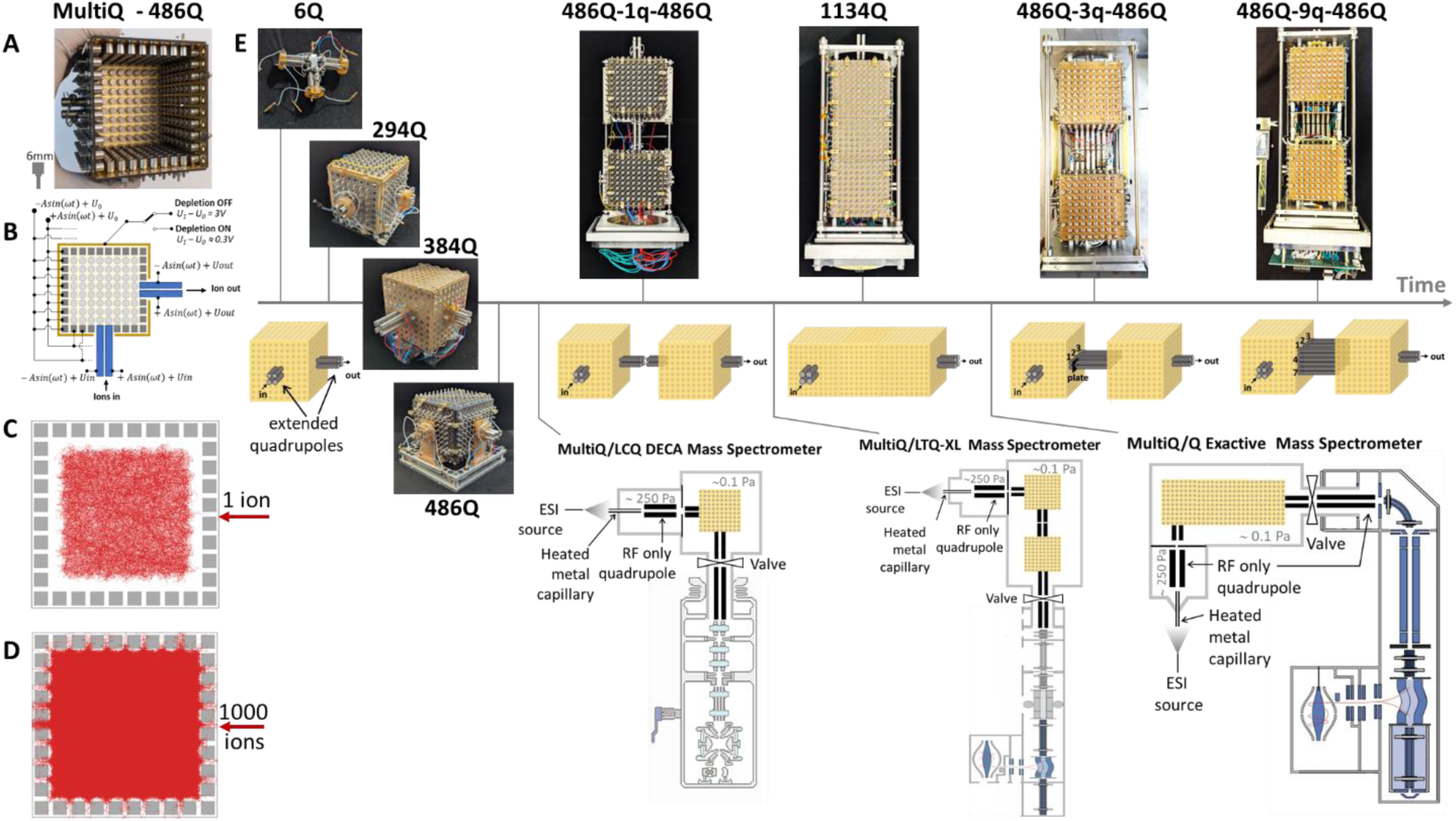
The MultiQ-Ion Trap. (**A**) An optimized 486-quadrupole (486Q) version of a MultiQ-IT with one wall removed for clarity. (**B**) Ions are trapped by the radiofrequency (RF) fields and a DC potential difference (ΔU =*Uwall - U0*). (**C**) Simulated trajectory of a single ion (mass 1500 Da, 3+ charge) projected onto a plane after being trapped with 150 V, 500 kHz RF, and ΔU = 4 V for 1 second. (**D**) Trajectories of 1000 of the same ions as in c but with the RF amplitude reduced to 50V and ΔU =0. (**E**) Development timeline of MultiQ-IT configurations with increasing numbers of quadrupoles (Q), from 6Q to 1134Q. The 486Q variant was further implemented in tandem configurations (486Q-1q-486Q, 486Q-3q-486Q, and 486Q-9q-486Q), with ion transfer enabled via extended quadrupoles (q). Example implementations into the different mass spectrometers (MS) used in the present studies are shown on the bottom right.

We first conducted a series of experiments that examined mixtures of peptide ions produced by an electrospray ionization source. These ions were introduced into a vacuum interface through a heated metal capillary (*21, 22*) and directed through an RF-only quadrupole ion guide to a single quadrupole port into the MultiQ-IT. Once trapped, the ions underwent thermalization through collisions with buffer gas molecules (N₂, 0.05–0.1 Pa) before exiting through one or more designated ports. The residence half-lives of ions within the trap ranged from 0.1 to 1 second, depending on specific ion characteristics such as molecular weight, charge state, and molecular composition (**figs. S3 and S4**). Direct ion current measurements demonstrated that the trap can transiently hold as many as 10^9^ – 10^10^ elementary charges per second (**fig. S5**). For ions with charge states ranging from +1 to +6, as determined from the mass spectral data, the maximum measured ion flux through the 486Q ion trap exceeded 10^9^ ions/s, which is 1000-fold higher than the capacity of ion traps used in state-of-the-art commercial mass spectrometers (*23–25*).

Coulombic repulsion between the large number of trapped ions with like charges can generate substantial electric fields – as high as 0.5–5 V/mm at the boundary of the ion cloud within the trap (**fig. S6**). Although the amplitude of the RF trapping signal could theoretically be increased to create a higher effective trapping potential to compensate for this Coulombic repulsion between ions, such an approach is only practical when the ions of interest are not required to explore the space between the quadrupoles. Therefore, to fully utilize the multiplicity of quadrupole outputs in the trap, the RF amplitude must remain low enough at a given RF frequency to maintain the Mathieu stability parameter of q<0.9; this constraint limits the strength of the effective trapping potential (*26*), in turn limiting the number of ions that can be practically confined in the MultiQ-IT, albeit in the present case to as many as 10^10^ elementary charges.

To evaluate the capability of the MultiQ-IT for manipulating ions in a massively parallel manner, we investigated the ability of the 486Q version to differentiate ions with distinct charge states in real time. Ions attempting to pass through any quadrupole port other than the designated entry and exit ports, can be effectively blocked by applying a sufficiently high potential difference between each quadrupole and its nearby wall (**Fig. 1B and figs. S1 and S2**). Because ions within the trap volume are largely thermalized, a potential difference of just 3V is sufficient to prevent all ions from exiting. As we reduce the potential difference towards 3/2k_B_T, singly charged ions begin to overcome the potential barrier, collide with the wall, and are lost. In contrast, thermalized multiply charged ions are less likely to penetrate the barrier, as its effective height increased proportionally with ion charge. Such enrichment of multiply charged ion species was previously observed using a single quadrupole ion trap, where ions (typically with charge states between +1 and +3) were first confined by raising a trapping potential barrier after injection. The barrier was then lowered to selectively deplete singly charged ions, followed by the release and detection of the remaining multiply charged species (*27, 28*). That approach required a sequential trapping and release mode of operation. In contrast, the MultiQ-IT enables simultaneous operation of a multitude of parallel ports for the selective removal of singly charged ions (in a manner somewhat analogous to on-line dialysis), greatly improving the efficiency and speed of charge separation. This property was confirmed both computationally (**Fig. 2A**) and experimentally (**Fig. 2B**) using several different mass spectrometers equipped with MultiQ-ITs (**Fig. 1E**). Indeed, by utilizing the full array of available ports concurrently, singly charged ions were depleted on a timescale comparable to their residence half-life within the trap. This allows real-time charge discrimination as ions transit the trap, eliminating the need for a sequential pulsing sequence.

**Fig. 2.**
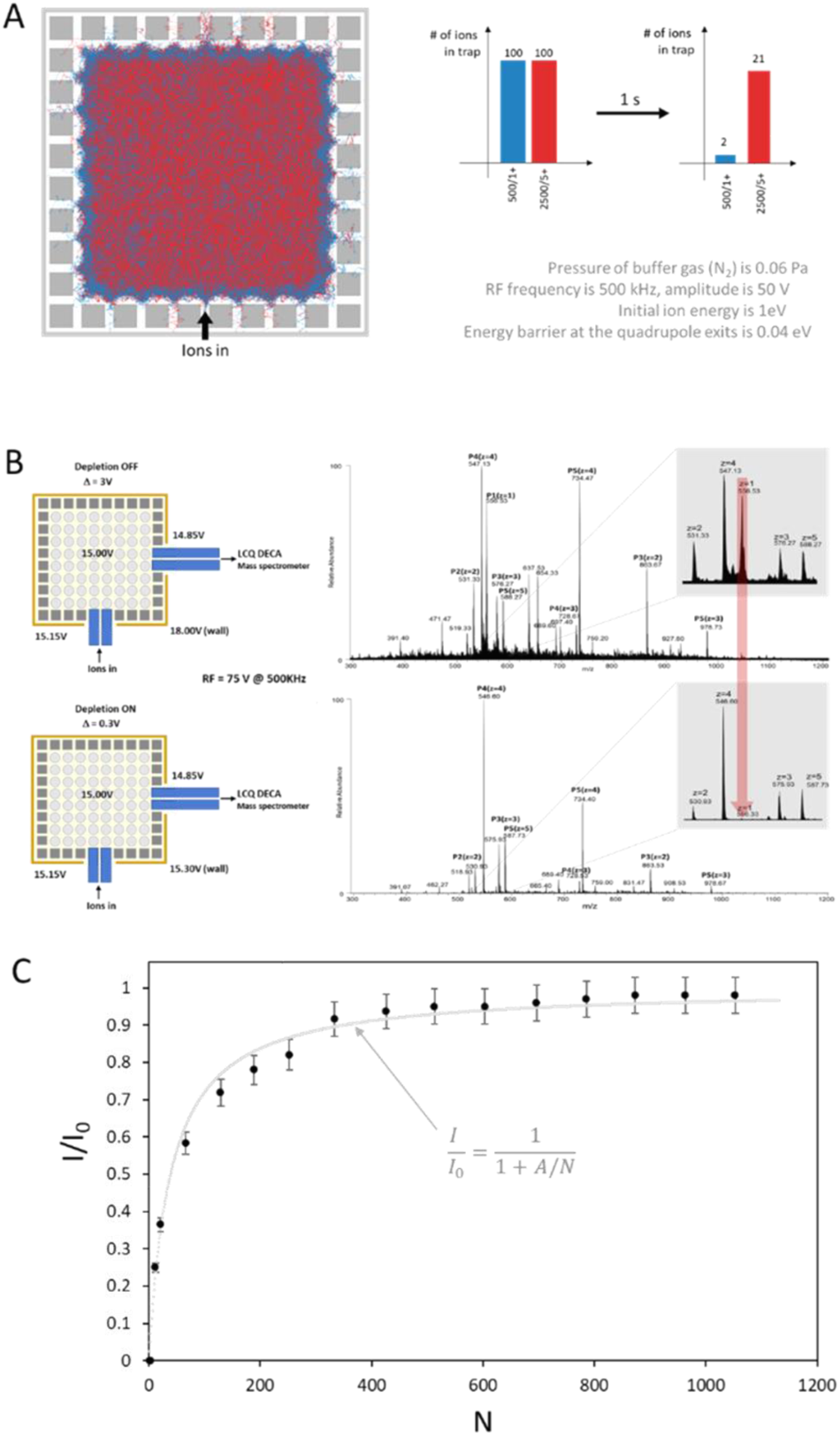
Selective depletion of singly charged ions using the MultiQ-IT. (**A**) Simulations of 200 ions in the 486Q MultiQ-IT using the Demiurge_MultiQ-IT_486 program (Supplementary). Trajectories are shown for 100 singly charged ions (m/z = 500; blue) and 100 multiply charged ions (m/z = 500, z = 5; red), projected onto a plane through the trap midpoint. Ions were tracked for up to 1 second, during which they could remain trapped, exit through a port, or collide with an electrode. (**B**) Experimental validation using a five-peptide mixture (800 amol/s) in a 486Q MultiQ-IT coupled to an LCQ DECA MS. With depletion OFF (top spectrum), singly charged ions dominate. With depletion ON (bottom), singly charged species are reduced, revealing multiply charged peptide signals. Left: potential distributions used in each mode. Peaks for peptides P1–P5 are labeled. (**C**) The number of output ports needed for efficient depletion was evaluated by plotting the normalized intensity (I/I₀) of singly charged ions as a function of available ports (N). Increasing N facilitates ion exit, reducing singly charged background in real time.

To determine the number of escape ports required for such efficient real-time depletion, we developed an expanded 1134-quadrupole version of the trap (**Fig. 1E**). This design, featuring a large number of ports, allowed us to systematically vary the number of exit ports with sufficiently low potential barriers to enable the preferential escape of singly charged ions from a mixed population of ions with charge states ranging from 1+ to 6+ (see, for example, **fig. S8, bottom panel**). Our experimental results (**Fig. 2C**) showed excellent agreement with a theoretical model based on the steady-state solution of Fick’s diffusion equation, which describes diffusion towards *N* circular absorbing apertures in a planar barrier (*29*). The quality of the fit was excellent (χ² = 0.0011 and R² = 0.999999), indicating that ion transport within the MultiQ-IT follows well-established principles of diffusion. For this specific geometry (**Fig. 1E**), we found that the singly charged ion outflow reaches half of its maximum value when the number of low-potential-barrier exit ports is *N* = 39 ± 3. This strong alignment between experiment and theory supports our hypothesis that ion motion and escape through multiple ports in the MultiQ-IT can be accurately modeled using the same theoretical framework that describes molecular diffusion to receptors on a cell membrane or transport to nuclear pore complexes on the periphery of the cell nucleus (*29*).

Next, we investigated whether the depletion process could be further enhanced by coupling two 486-quadrupole traps in tandem, each designed to reduce the population of singly charged ions by a factor of ∼10 while at the same time efficiently trapping higher-charge-state ions. Since the two traps function independently, their effect in tandem should be multiplicative. As predicted, this approach resulted in a ∼100-fold improvement in the signal-to-noise ratio for a low concentration five-peptide test mixture after selectively depleting singly charged ions from the primary ion beam (**Fig. 3**). Prior to depletion, peaks corresponding to only two peptides were faintly discernable amid a dense background of singly charged ion peaks associated with the so-called chemical noise (*30*) (**Fig. 3A**). After depletion, however, all peptides in the sample became clearly observable (**Fig. 3B**). A similar striking effect is seen in **Figs. 3C and 3D**, where spectra recorded before and after depleting a dominant singly charged contaminant are compared, revealing the previously obscured low intensity protein signal.

**Fig. 3.**
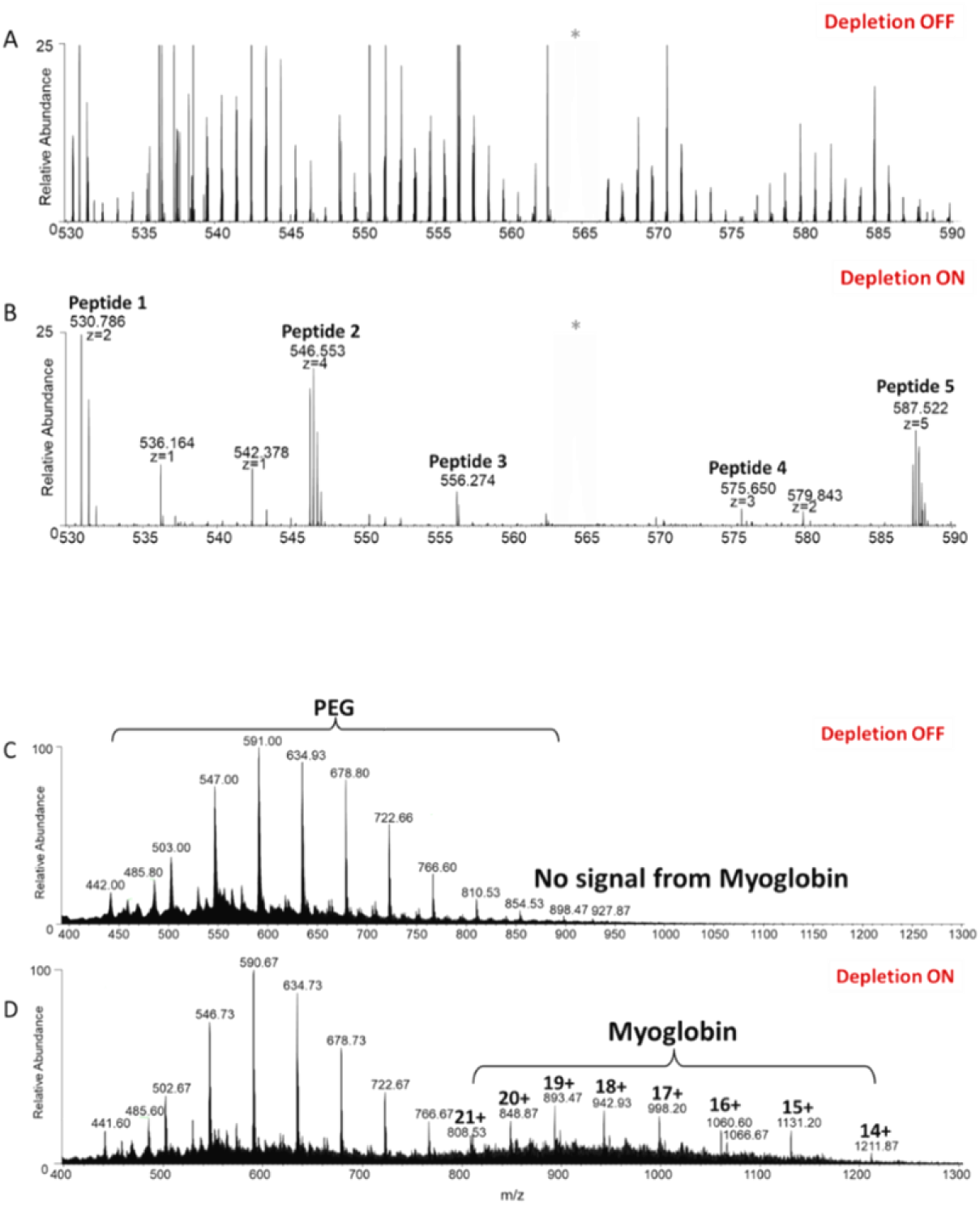
Signal-to-noise enhancement via selective depletion of singly charged ion background. Portion of the mass spectra arising from a five-peptide mixture electrosprayed at 17 amol/s, recorded without (**A**) and with (**B**) selective depletion of singly charged ions. In the absence of depletion (**A**), peptide signals are masked by “chemical noise” from abundant singly charged background ions. When depletion mode is enabled (**B**), using the tandem 486Q MultiQ-IT trap (Fig. 1), singly charged ions are removed in real time as ions transit the trap system. This results in a 100-fold reduction in background, revealing clear signals for all five peptides. Asterisks indicate artifact peaks arising from pickup events in the LTQ-XL mass spectrometer used in this experiment. (Mass spectra of myoglobin electrosprayed at 17 amol/s in the presence of a 100-fold molar excess of polyethylene glycol (PEG 500; a common contaminant in protein preparations), recorded without (**C**) and with (**D**) singly charged ion depletion. (**C**) Depletion off – no signal from myoglobin is observed. (**D**) Depletion on – singly charged ions from PEG 500 are preferentially removed, allowing for the observation of the multiply charged myoglobin ion peaks.

**figs. S7–S9** provide additional examples demonstrating the efficacy of the MultiQ-IT in depleting singly charged species across various instrument configurations. In all these cases, we observe great improvements in the signal-to-noise ratios. We also explored the possibility of selectively depleting intermediate charged states (e.g., 2+ and 3+), in addition to 1+, as a strategy to enhance the detection of species with charge states of 4+ or higher (**figs. S10 and S11**) This approach, which we term “extended charge state depletion”, involves appropriately lowering the effective potential barrier to enable the selective escape of 2+ and 3+ ions along with singly charged species. The resulting increase in the relative abundance of higher charge state ions greatly improves the detection of extremely low-level analytes (**fig. S12**) and holds considerable promise for enhancing the identification of chemically crosslinked species in mass spectrometric workflows (*31, 32*) (**fig. S13**), which has become a powerful approach to assist in the integrative structural modeling of large protein-containing assemblies (*33–36*). However, to fully unlock the transformative potential of this capability for detecting trace, highly charged species, it will be necessary to implement a truly parallel mass spectrometric system, i.e., one that comprises a large array of dedicated, fragmentation-capable m/z analyzers operating simultaneously, with each channel continuously acquiring data throughout the entire temporal profile of even the weakest signals. Realizing such a system, catalyzed by the development of the MultiQ-IT, represents an ambitious – yet, we believe, achievable – goal, one we hope to see fulfilled in the near future.

As an initial step toward this goal, we sought to demonstrate how the MultiQ-IT can support large-scale parallel ion manipulation using its existing architecture. As shown above, applying a modest potential difference across the exits of the MultiQ provides a simple yet effective demonstration of how multiple outputs can be harnessed, without requiring any structural modifications to the original MultiQ-IT design. However, our broader objective is to demonstrate how true massive parallelism in mass spectrometry can be realized by incorporating more advanced ion transport strategies through these outputs – specifically, those capable of real-time beam splitting into multiple, narrow m/z-range sub-beams for concurrent detection or analysis. In a first step to explore this concept, we employed a tandem MultiQ-IT configuration to evaluate whether multiple output ports could function as independent m/z filters without requiring a seperate mass analyzer at each port. In this configuration, the first MultiQ-IT operates as an ion beam splitter, while the second functions as a beam combiner, allowing the output of different channels to be assessed collectively using a single mass analyzer. We began by connecting two traps using three extended quadrupoles and subsequently scaled up to nine, creating systems with three and nine parallel channels, respectively (**Fig. 1E**). To selectively transmit ions within defined *m/z* windows, we tested various configurations of the extended quadrupoles, examining their performance as conventional mass filters and as linear traps with axial ion ejection (*18–20, 37*). Unlike prior designs, our approach sought to efficiently return non-resonant ions (those outside a given m/z filter window) back to the interior of the MultiQ-IT, allowing them to further explore and pass through alternative filters better suited to their m/z. Through iterative modeling and prototype testing, we optimized one such design that featured nine extended quadrupoles configured for resonant ion ejection (**Fig. 4A**). Our modeling included detailed computer simulations of ion motion within a single quadrupole channel (**Fig. 4B**). These simulations demonstrate efficient and selective ion transmission for target *m/z* values while maximizing ejection efficiency and minimizing ion losses. The quadrupoles are short (1.2 cm), with two adjacent rods tilted slightly (∼0.01 rad) along one axis and parallel along the orthogonal axis – an arrangement we term the “tilted quadrupole” (**Fig. 4A**). A small excitation voltage applied across the non-inclined rods induces resonant excitation of ions within a specific m/z range, gradually increasing their oscillation amplitude. Once sufficiently displaced, ions encounter the flared potential created by the inclined rods, which accelerate them toward the exit, enabling them to overcome a small stopping potential and leave the MultiQ-IT. Our simulations (**Fig. 4C**) indicate that this approach should achieve a resolving power of 50 (*Δ* = 10 at *m/z* = 500), an 80% ejection efficiency at resonance with minimal losses due to collisions with the electrode, and 100% efficiency for return of non-resonant ions to the MultiQ-IT ion confinement region.

**Fig. 4.**
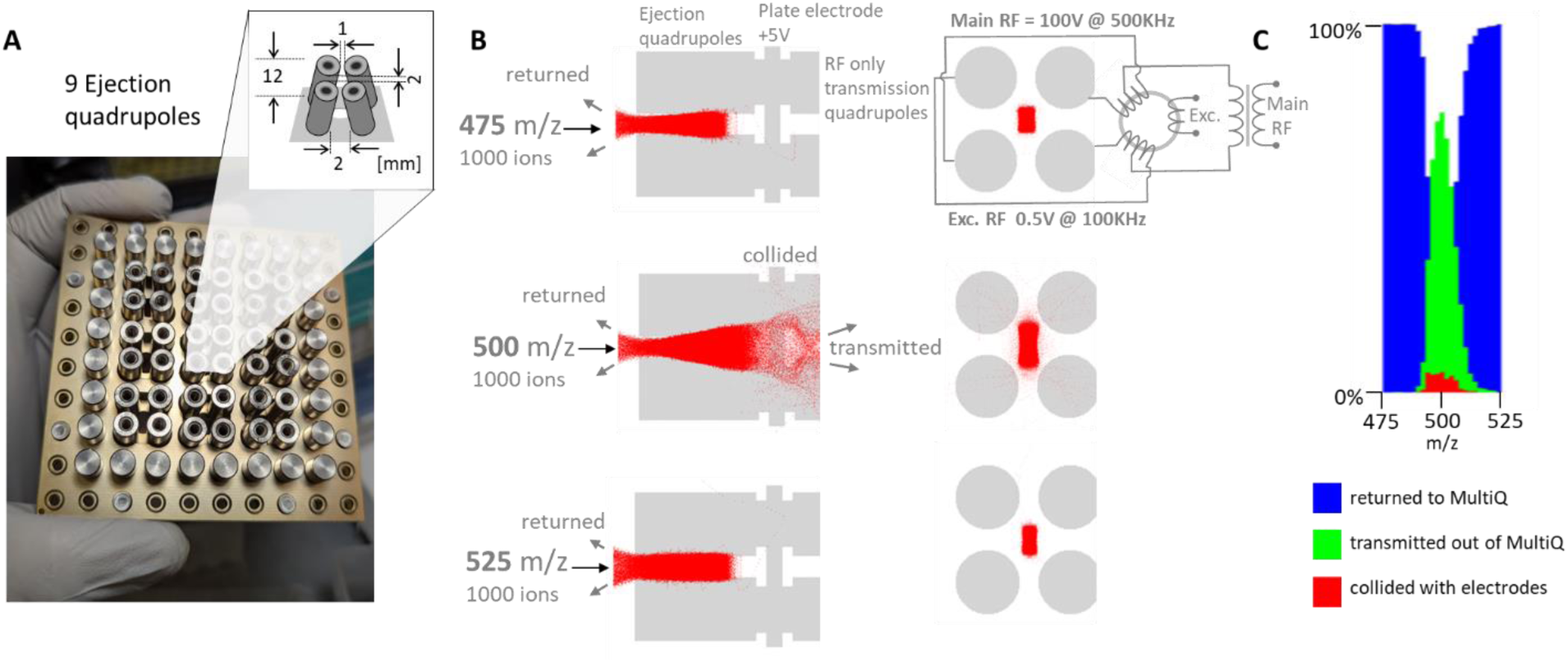
MultiQ-IT system for continuous ejection of ions into multiple m/z-specific sub-beams. (**A**) Section of the 486Q MultiQ-IT showing 9 tilted ejection quadrupoles integrated into the ion trap wall. The construction and orientation of these quadrupoles enable selective ion transmission. (**B**) Simulations of 1000 ions with m/z 475, 500, and 525 entering the tilted quadrupoles. An auxiliary excitation RF is applied to selectively transmit ions centered around m/z 500, as indicated. (**C**) Simulation results showing that ions with m/z values significantly lower or higher than 500 are returned back into the MultiQ-IT, while ions centered near m/z 500 are preferentially transmitted. A minor fraction of partially transmitted ions is lost due to collisions with the quadrupole electrodes.

Experimental validation of the computationally predicted operation mode of the tandem MultiQ-IT, with two ion traps connected via nine parallel channels (**Fig. 1E, top right**), is presented in **Fig. 5**. Ubiquitin ions were selectively transmitted through four of these channels, each tuned to continuously pass ubiquitin ions with a distinct charge state. The system successfully split the incoming ion beam into four distinct sub-beams (**Fig. 5A**), each centered on a selected m/z value and confined within a ±50 m/z unit window. These sub-beams were subsequently recombined in the second MultiQ-IT, yielding a composite mass spectrum from all four channels (**Fig. 5B**). For comparison, **Fig. 5C** shows the control spectrum obtained by transmitting ubiquitin ions directly from the first to the second trap under a DC gradient, without applying resonant RF excitation – i.e., without m/z-selective splitting. The measured ion ejection efficiency in the parallel splitting mode was approximately 10% of that observed during direct, unsplit transmission. While these experimental results qualitatively confirmed the core predictions of our computational model, deviations in ion ejection efficiency and m/z resolution are noted. These discrepancies are likely attributable to minor misalignments in rod inclination angles and overall quadrupole geometry, which are areas targeted for refinement in future device iterations. We also envision several alternative designs for achieving m/z-selective beam separation with enhanced resolution and improved ion transmission efficiency.

**Fig. 5.**
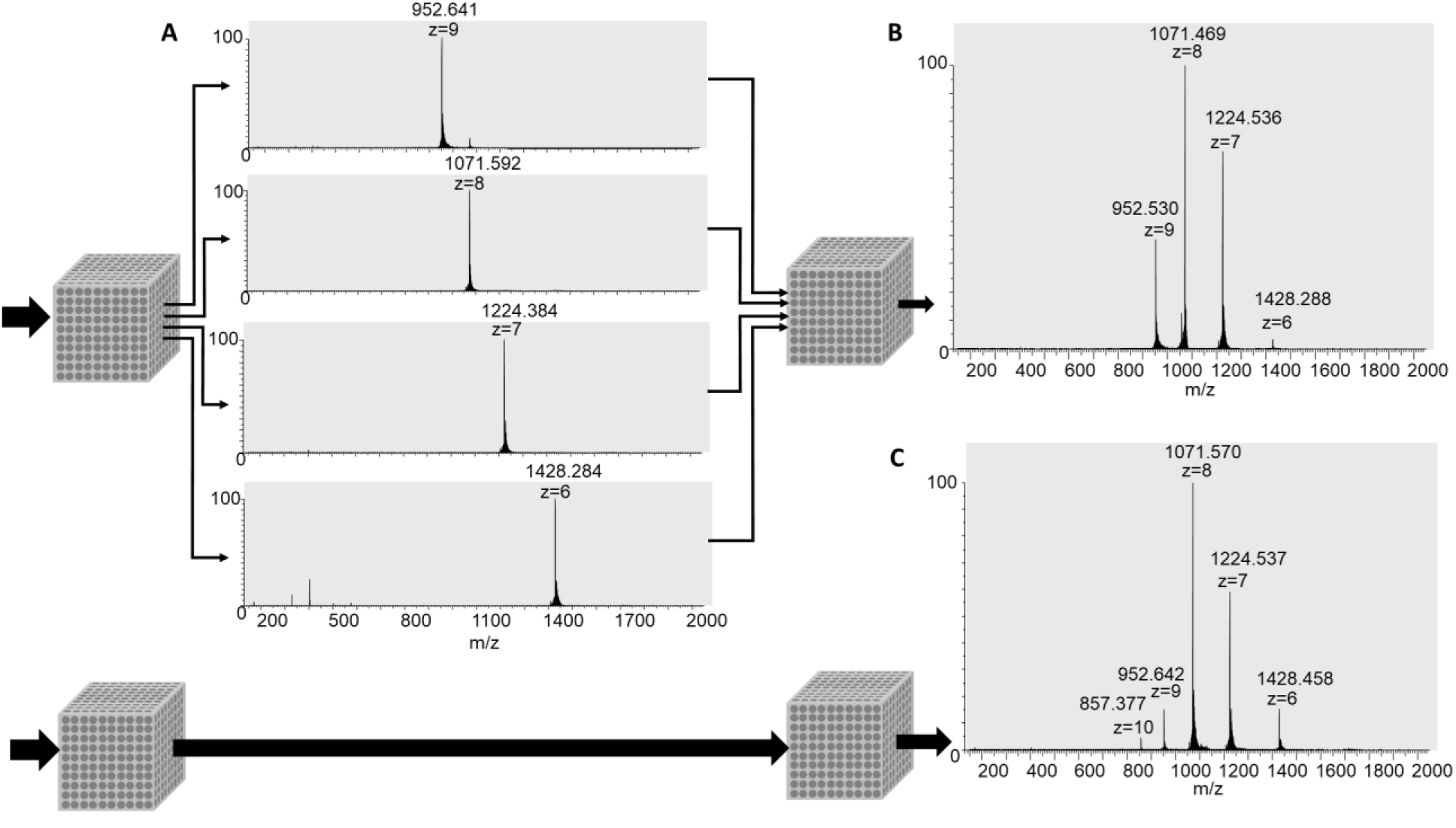
Splitting an ion beam into parallel sub-beams. Four randomly selected channels from the nine available in the tandem MultiQ-IT system were independently optimized to transmit distinct ubiquitin charge states by tuning the resonance frequency (40–60 kHz) and RF amplitude (ranging from 1.5–5 V) applied to the extended quadrupoles. Once optimized, ion transfer between the first and second MultiQ-IT traps could be selectively blocked by applying a reverse DC gradient. (**A**) m/z spectra recorded by selectively activating each channel, allowing isolation and detection of individual charge states. (**B**) Simultaneous activation of all four optimized channels resulted in combined transmission of selected ions into the second MultiQ-IT, generating a composite ubiquitin spectrum. (**C**) Control spectrum acquired by transmitting ubiquitin ions through a single channel without applying any resonant excitation, using only a DC gradient. The observed ion ejection resolution was ∼10 (theoretical ∼50). Ubiquitin was chosen for its well-resolved charge states (+6 to +10), making it an ideal model for assessing multi-channel selectivity.

In this work, we have demonstrated that the MultiQ-IT platform holds significant promise for enabling massively parallel mass spectrometry, offering a transformative strategy to enhance sensitivity, speed, and dynamic range in MS analyses. Unlike conventional ion traps that operate with a single input and output, requiring sequential ion accumulation and ejection, the MultiQ-IT incorporates multiple quadrupole-based inputs and outputs. These serve two key roles: (i) transient containment and thermalization of ions, as evidenced by our ability to trap and manipulate up to 10^9^ ions, exceeding the capacity of existing designs by three orders of magnitude; and (ii) controlled ion release in which ions, driven by entropy can distribute across all available exits, so forming spatially separated sub-beams, whose composition can be modulated to optimize downstream analysis.

This ability to generate and manipulate spatially separated ion sub-beams is foundational for achieving true massive parallelization in mass spectrometry. Our findings establish the feasibility of parallel mass spectrometry using the MultiQ-IT. Much like nucleocytoplasmic transport to and through nuclear pore complexes (NPCs), ion movement in the MultiQ-IT is governed by diffusion-driven transport, regulated by the availability and configuration of input and exit ports. In our charge state depletion studies, we found that effective ion depletion requires only a modest number of quadrupole ports – much like NPCs, which efficiently mediate molecular exchange despite their limited number (*38*). Using the steady-state solution to Fick’s diffusion equation, we determined that a few hundred ports are more than sufficient to achieve optimal ion depletion, mirroring the efficiency of NPCs in biological systems (*29, 38*). The MultiQ-IT is also highly scalable, following an N² relationship (specifically, 6(N−1)² for a cubic configuration, where *N* represents the number of cylindrical electrodes along a single dimension). This scalable architecture presents compelling opportunities in proteomics, metabolomics, and single-cell analysis, particularly when each sub-beam is coupled with integrated fragmentation and spectrum acquisition capabilities. Recent advances in miniaturized and array-based mass spectrometer systems (*39*) suggest that such capabilities are increasingly within technological reach. Indeed, the development of the MultiQ-IT is expected to further accelerate these technological advances by providing a robust platform ideally suited for such integration The MultiQ-IT represents a significant advance towards parallel mass spectrometry, offering new avenues to improve speed, sensitivity, dynamic range, redundancy, and system scalability. Just as parallel computing transformed information processing, the MultiQ-IT has the potential to redefine what is possible in analytical science, opening the door to next-generation applications in chemistry, biology, and medicine.

## Acknowledgements

**Funding:** This work was supported by a NIH Director’s Transformative Research Award to BTC (PHS R01GM136654).

**Author Contributions:** All aspects of this work were contributed to by both authors.

**Additional Contributions:** We would like to thank Herbert Cohen for assistance with electronics during the early stages of this work. We also thank Stephen Kent and Michael Rout for reading the draft manuscript and providing us with feedback and encouragement.

**Competing Interests:** The authors declare that they have no competing interests.

**Data and materials availability:** All processed data are available in the main text or the supplementary materials. All raw data is available upon request. Simulation code is available at: https://github.com/akrutchins.

## Supplementary Materials

### Materials and Methods

#### MultiQ-IT Instrumentation

Details of the construction and setup of the MultiQ-IT ion trap are shown in Figures S1 and S2. Ion currents were monitored using a low-noise current amplifier (Stanford Research Systems, Model SR570). Pulse generation and resonant excitation of ions were carried out using digital signal generators (Stanford Research Systems Models DS345 and DG535) and a Koolertron DS Signal Generator/Counter.

#### Reagents and Sample Preparation

A mixture of five synthetic peptides (AnaSpec) was prepared at concentrations ranging from 1 to 100 fmol/μL in 0.01% formic acid in 70:30 (v/v) water/acetonitrile. The peptide mixture included:

– [Leu⁵]–Enkephalin (YGGFL, 555.269 Da; AS-24333)
– [Des-Pro²]–Bradykinin (RPPGFSPFR, 1059.561 Da; AS-20667)
– Bak BH3 (GQVGRQLAIIGDDINR, 1723.933 Da; AS-61616)
– Neuropeptide S, mouse (SFRNGVGSGAKKTSFRRAKQ, 2181.188 Da; AS-61246)
- ACTH (1–24), human (SYSMEHFRWGKPVGKKRRPVKVYP, 2931.580 Da; AS-20613)

This mixture generated distinct ESI peaks in the 500–600 m/z range, with ion charge states from +1 to +5, enabling detailed evaluation of depletion and trapping performance. Other analytes used included ubiquitin (bovine erythrocytes, Sigma-Aldrich U6253), myoglobin (equine heart, Sigma-Aldrich M1882), and polyethylene glycol (PEG 350, Fluka 81318), prepared at 0.1–1 pmol/μL in the same solvent.

#### Software and Data Analysis

Mass spectra were visualized using Xcalibur and FreeStyle (Thermo Fisher Scientific). Spectra are presented unprocessed unless otherwise noted. Ion motion within the MultiQ-IT was simulated using custom Processing and Java-based software, based on the model described by Krutchinsky et al. (J. Am. Soc. Mass Spectrom. 1998, 9, 569–579). Simulation code is available at: https://github.com/akrutchins.

## Supplementary Figures

**Fig. S1.**
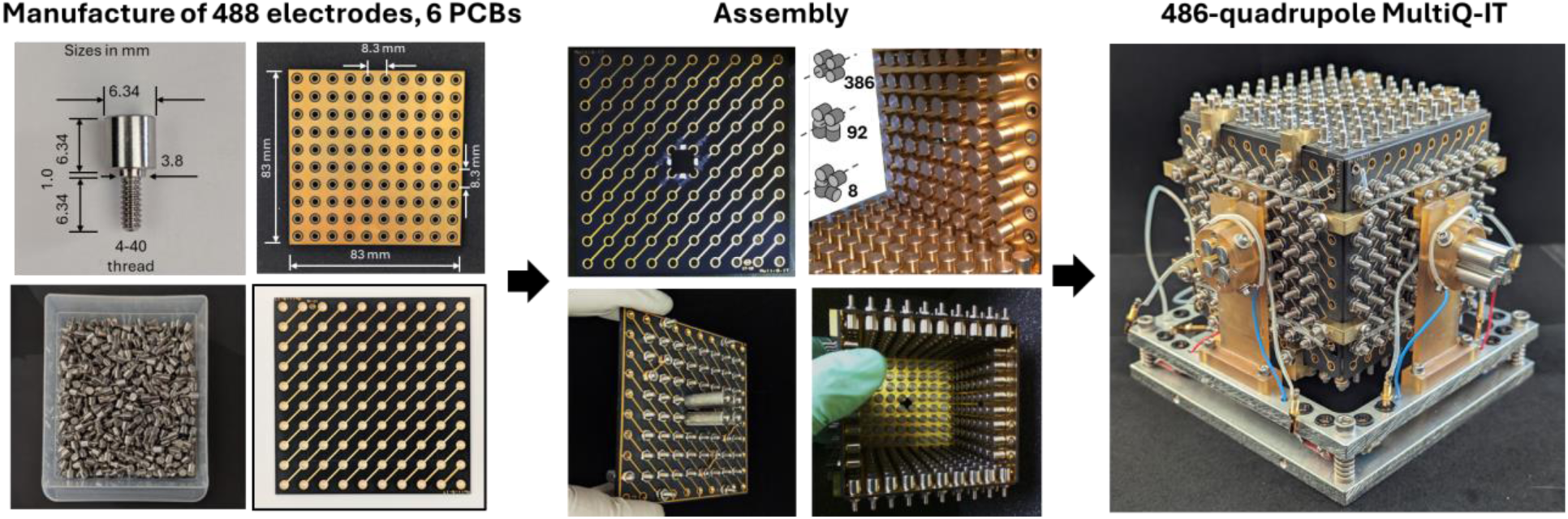
Manufacture and assembly of the 488-electrode (486-quadrupole) version of the MultiQ-IT. Manufacture: Thousands of quadrupole electrodes were custom-manufactured by Sandum Precision Industry Co., Ltd. (China), then washed and assembled onto six printed circuit boards (PCBs) custom-made by JLCPCB (Hong Kong, China). **Assembly:** The electrodes were mounted on the printed circuit boards, which were then assembled into a “cube” to form the ion confinement region, with the quadrupole electrodes facing inward. Two PCB plates contained 100 electrodes each, two had 80 electrodes each, and the remaining two contained 64 electrodes each, ensuring continuity of the quadrupole array at the cube’s corners. This configuration produced three types of quadrupoles, as shown in the figure inset. The trap comprises 486 quadrupoles: 386 “classical” quadrupoles with cylindrical symmetry axes aligned with their quadrupole axes, and 100 “non-classical” quadrupoles with cylindrical symmetry axes that do not fully align. Simulations and experiments indicate that the “non-classical” quadrupoles, due to their fringing fields and smaller effective RF field radii, may improve ion confinement while maintaining efficient ion transmission under permissive conditions (see Fig.1d in the main text). Some PCBs include holes to facilitate ion passage through extended quadrupoles that are positioned on the outer face of the cube (middle figure). The steps illustrated here were also used to construct the other versions of the MultiQ-IT alluded to in the main article. **486-quadrupole MultiQ-IT:** A fully assembled version of the trap is shown on the right.

**Fig. S2.**
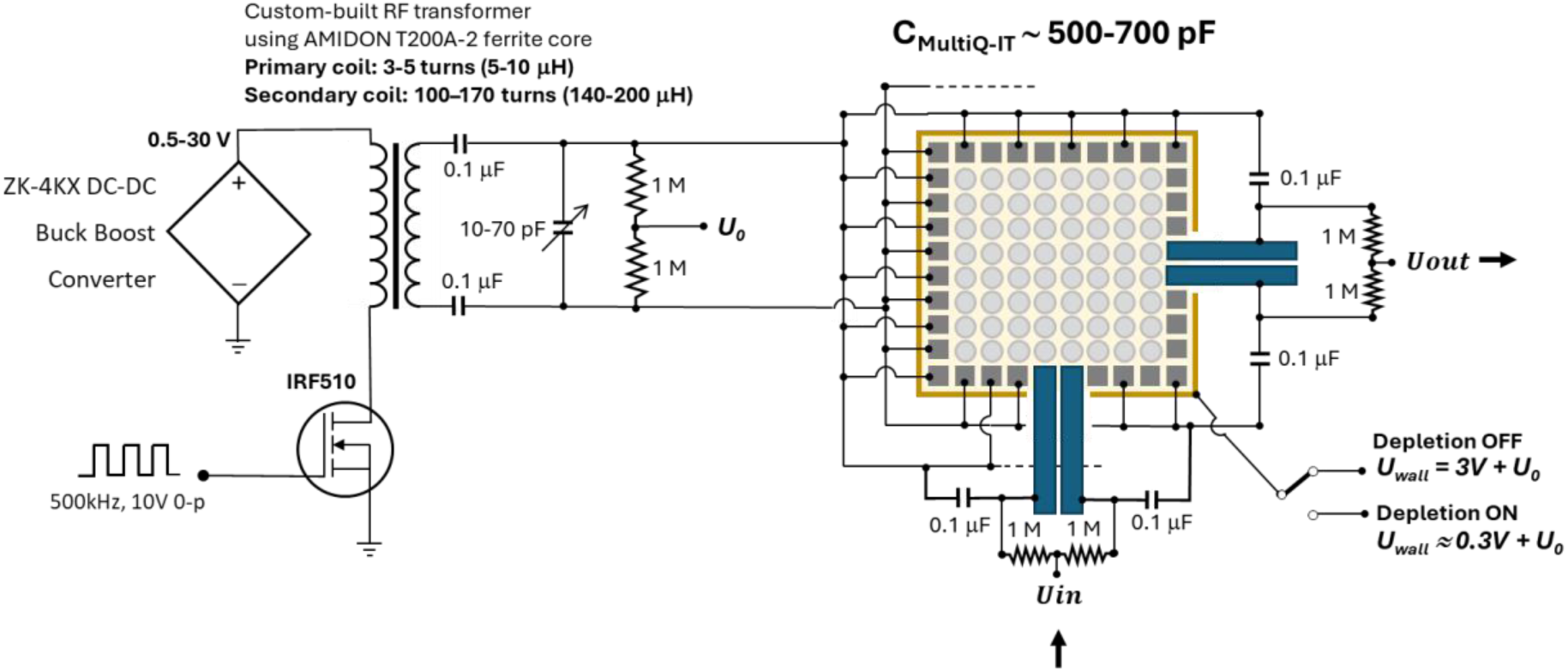
Schematic diagram of a custom RF power supply driving a single MultiQ-IT. The number of turns in the secondary coil of the custom-wound RF transformer was selected to match the resonance range of the LC circuit, where the major contribution to C comes from the capacitance between the connected quadrupole rods of the trap. Fine tuning of the resonance is achieved using a variable 10–70 pF capacitor (Jameco Valuepro CTC08-70-R). Various other simple circuits were used to control ion transport through one or more extended output quadrupoles; the specifics of these circuits are indicated in the relevant figures.

**Fig. S3.**
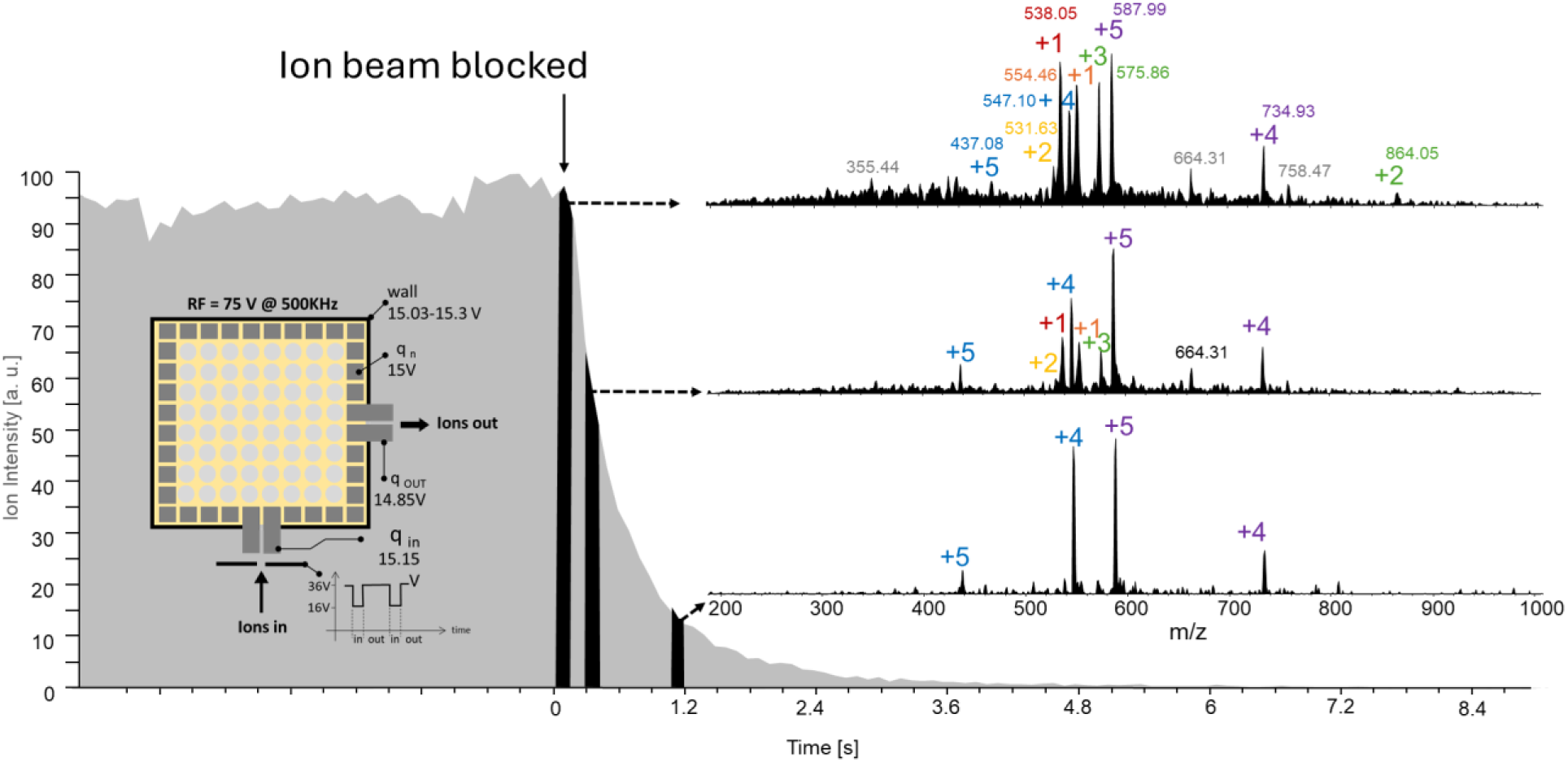
Measurement of ion residence times inside the 486-quadrupole MultiQ-IT. A continuous ion beam from an ESI ion source was periodically blocked by raising a potential applied to an electrode positioned upstream of the entrance quadrupole of the MultiQ-IT, during which time ions exiting the trap were continuously sampled using an LCQ DECA mass spectrometer. The exponential decay of the total measured ion intensity yielded a half-life of ∼0.3 ± 0.1 s. Notably, ions of higher molecular mass with higher charge states, +4 and +5, tend to remain in the trap longer than do ions with lower charge states. The electrospray solution of the mixture of peptides was introduced at a flow rate of 1.7fm/s.

**Fig. S4.**
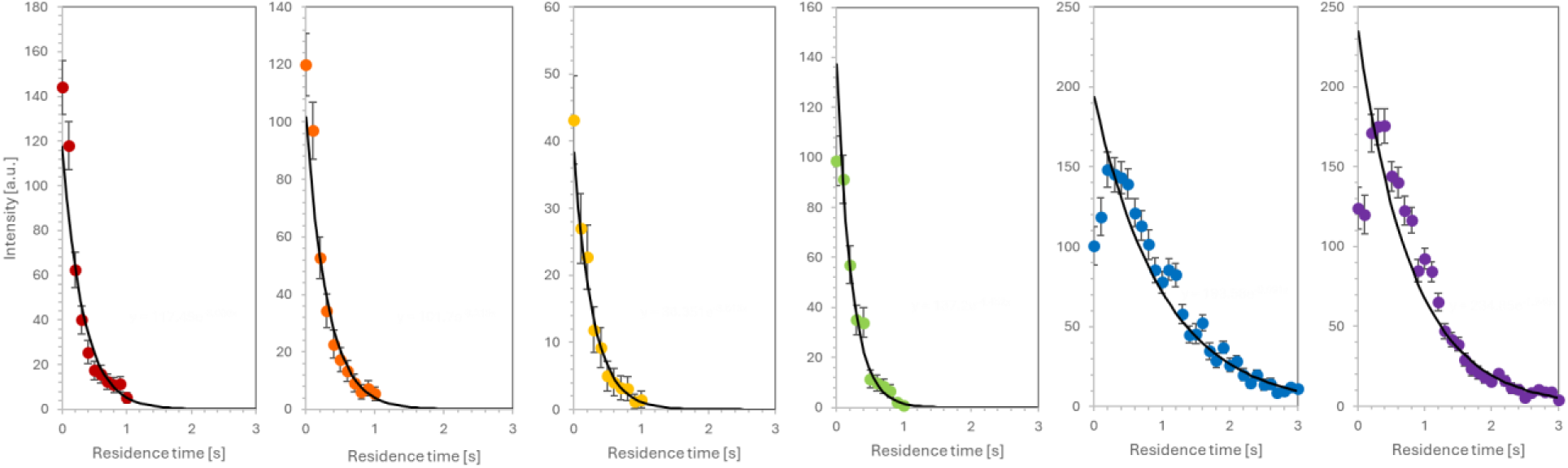
The ion residence half-lives within the 486-quadrupole MultiQ-IT of various peptide ions. These were measured by the decrease in ion intensity exiting the trap after the ion beam entering the trap was blocked (see fig. S3).

**Fig. S5.**
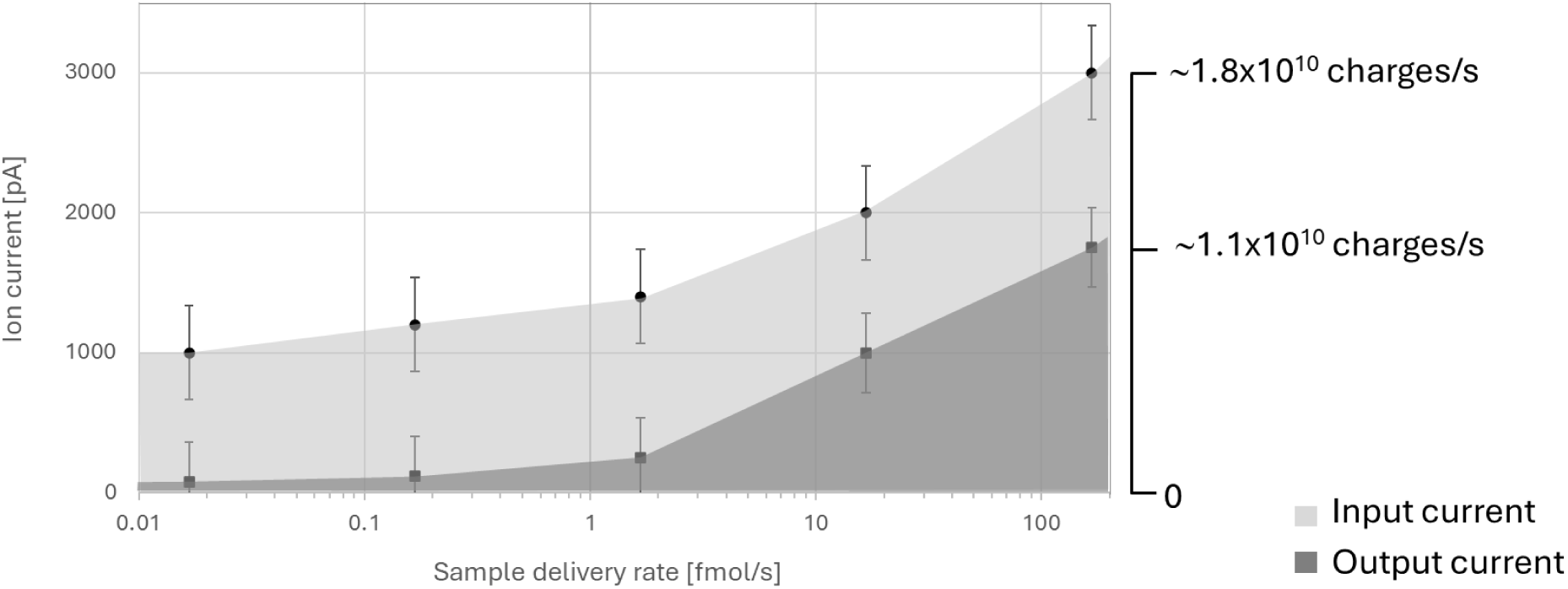
Experimentally determined ion capacity of the 486-quadrupole MultiQ-IT. Top curve. Total ion current generated by an ion source electrospraying a 5-peptide mixture as a function of concentration at a flow rate of 17 nL/s (light grey). **Bottom Curve.** The current of ions trapped for periods ∼1 sec (see figs. S3 and S4) before exiting through a single output is shown in dark grey. The ion currents were measured using a low-noise current preamplifier (Stanford Research Systems, Model SR570). The 486-quadrupole MultiQ-IT demonstrated the ability to transiently trap as many as ∼10^10^ elementary charges per second for high concentrations of the electrosprayed sample.

**Fig. S6.**
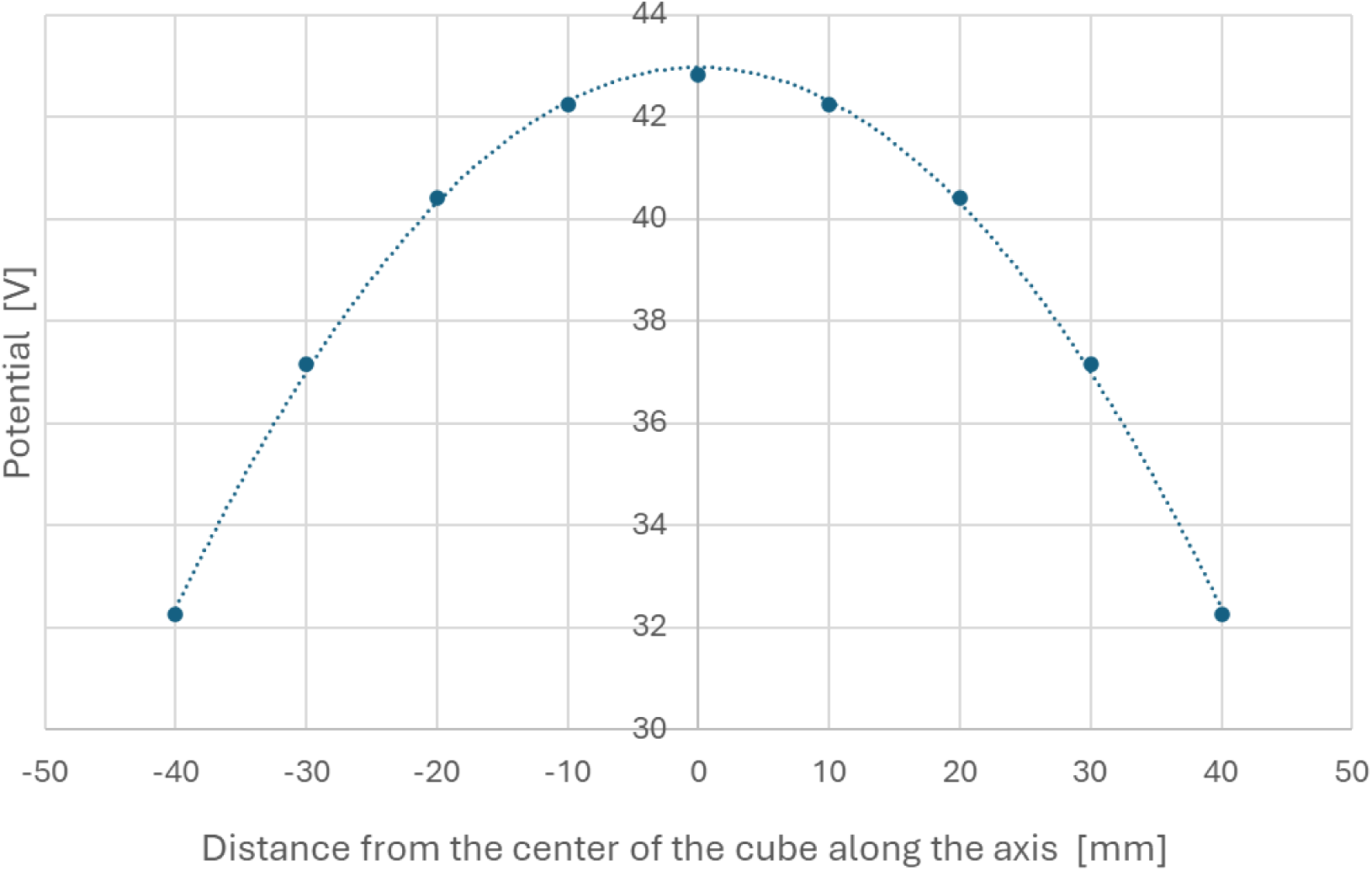
Electric potential across a cubic volume containing 10^9^ randomly positioned elementary charges. The cube dimensions are 80 mm × 80 mm × 80 mm – i.e., the size of the inner volume of the 486-quadrupole MultiQ-IT. Potentials were calculated (https://github.com/akrutchins/Coulomb-CUBE) at nine points spaced 10 mm apart along an axis passing through the center of one face. This data provides an estimate of the electric field created by Coulomb repulsion at the boundary of the ion cloud inside the MultiQ-IT, where the electric field is estimated to be approximately 0.5 V/mm.

**Fig. S7.**
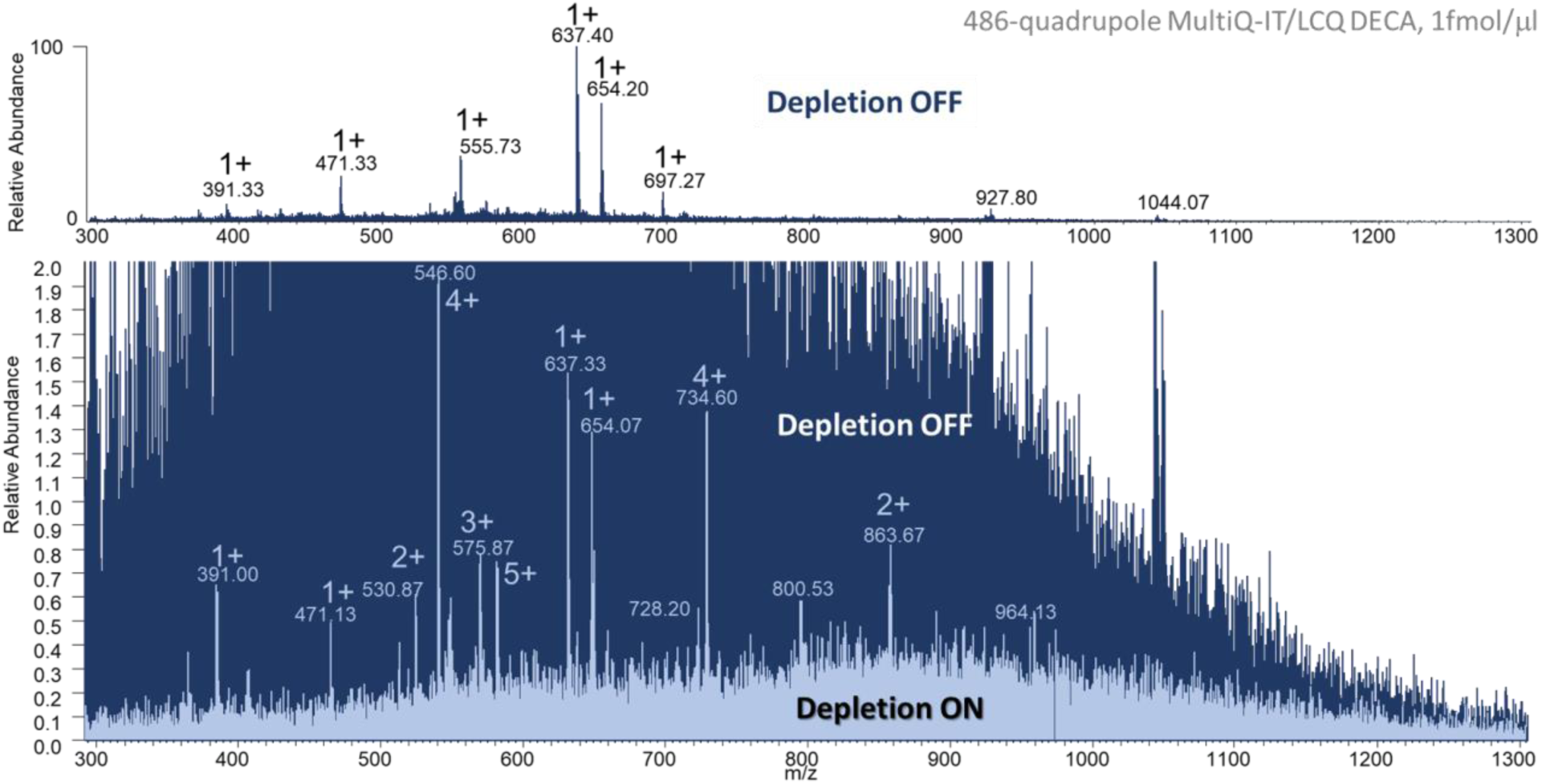
Selective depletion of singly charged ions in the tandem MultiQ-IT/LTQ-XL mass spectrometer. ESI spectrum of a mixture of 5 peptides directly infused at a rate of 8 amole/s into a 486-quadrupole MultiQ-IT/LCQ DECA mass spectrometer. **Upper panel**: Mass spectrum with Depletion mode OFF. The spectrum is observed to be dominated by singly charged species. **Lower panel:** The upper spectrum (dark blue) was obtained with Depletion mode OFF (50-fold vertical expansion of the spectrum in the upper panel, while the lower spectrum (light blue) was obtained with the MultiQ-IT set to Depletion mode ON (see settings in Extended Data Fig.8C) with no vertical expansion. The spectra are overlaid to highlight the extent of singly charged ion removal, enabling clear observation of signals from all peptides with an approximately 70-fold improvement in the signal-to-noise ratio.

**Fig. S8-Part I.**
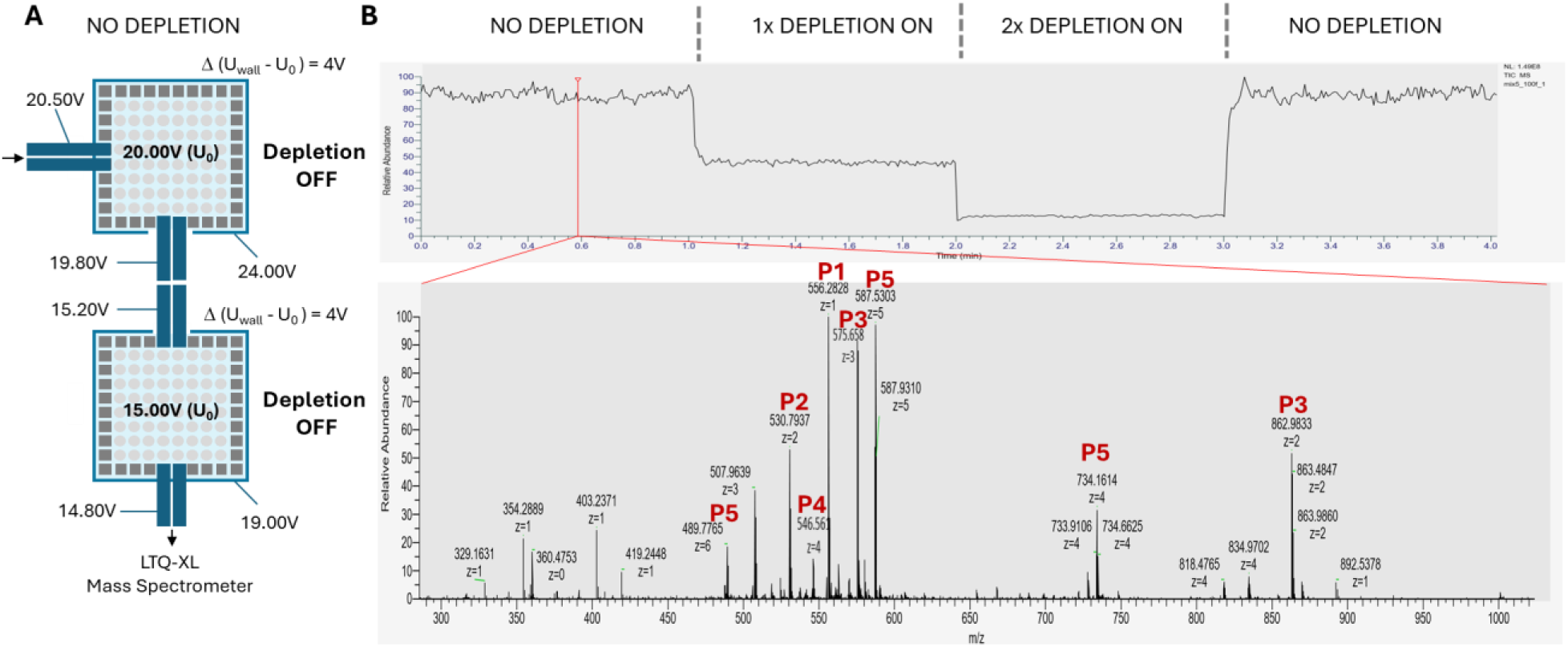
Depletion OFF control for singly charged ion depletion in the tandem MultiQ-IT/LTQ-XL mass spectrometer. An equimolar mixture of 5 peptides was electrosprayed at a flow rate of 1.7 fmol/s. (**A**) Distribution of potentials in the tandem 486-quadrupole MultiQ-ITs in NO DEPLETION mode. (**B**) (Top) Total ion count (TIC) and (bottom) a single-scan m/z spectrum in NO DEPLETION mode. The major peaks arising from the 5-peptide mixture are labeled in the spectrum. Note that the relative intensity of the singly charged peptide signal at m/z = 556.282 (labeled as P1) is 100 when the tandem MultiQ-IT is in NO DEPLETION mode, Δ(U_wall_-U_o_)=4V, where U_wall_ is the voltage applied to the MultiQ-IT wall and U_o_ is the DC voltage offset applied to the quadrupoles. The RF voltage was applied to the quadrupoles as a sign wave with an amplitude equal to 75V and a frequency of 500 kHz. This same RF voltage was applied to the MultiQ-ITs used to obtain the data presented in all the subsequent Supplementary Figures.

**Fig. S8-Part II.**
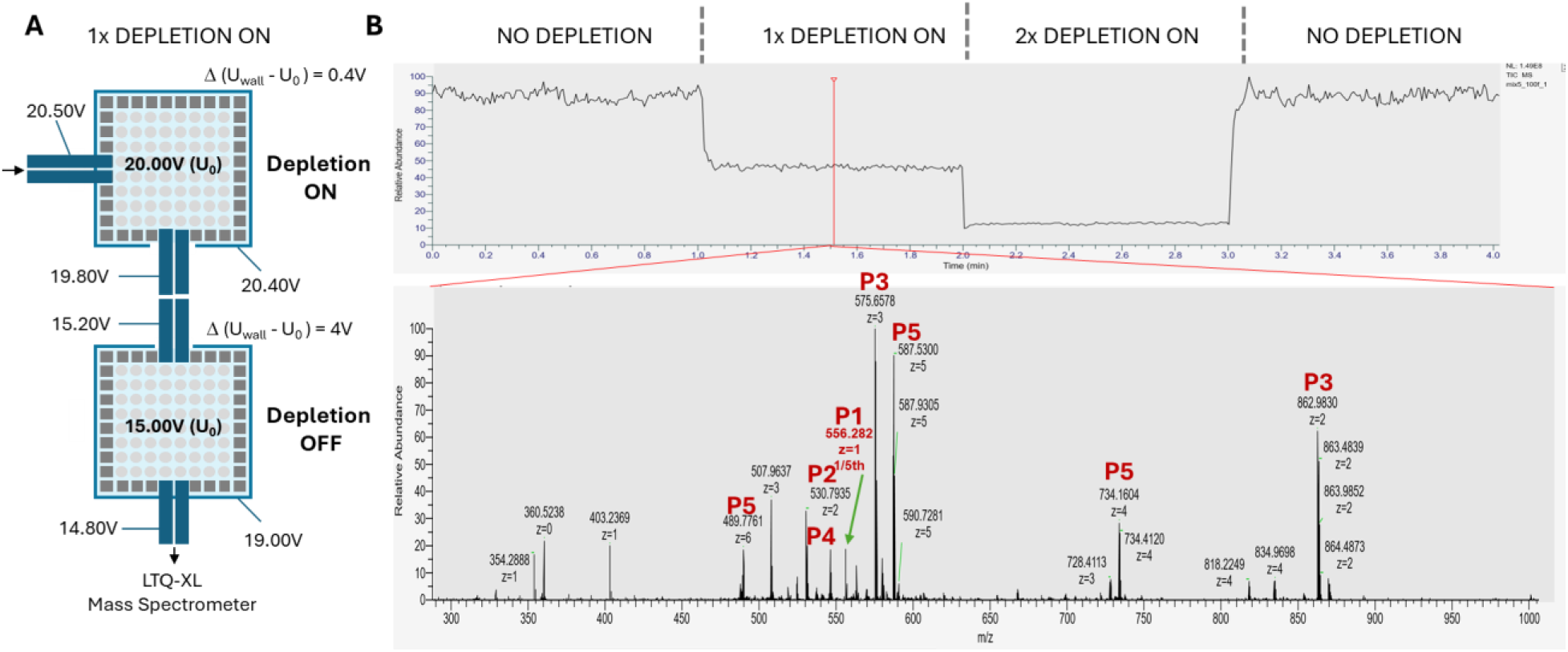
Selective depletion using only the first trap of the tandem MultiQ-IT/LTQ-XL mass spectrometer. An equimolar mixture of 5 peptides was electrosprayed at a flow rate of 1.7 fmol/s. (**A**) Distribution of potentials in the tandem 486-quadrupole MultiQ-ITs in 1x DEPLETION mode (1st MultiQ in Depletion ON mode, 2nd MultiQ in Depletion OFF mode) where the difference between the DC offset of the first trap quadrupoles and the wall was reduced to 0.4 V. (**B**) Total ion count (TIC) and a single-scan m/z spectrum in 1x DEPLETION ON mode. All major peaks of the 5-peptide mixture are labeled in the spectrum. Note that the intensity of the singly charged peptide signal at m/z = 556.282 (labeled P1) is reduced by a factor of 5 in this mode compared with the signal in Depletion OFF mode (fig. S8B-part I above).

**Fig. S8-Part III.**
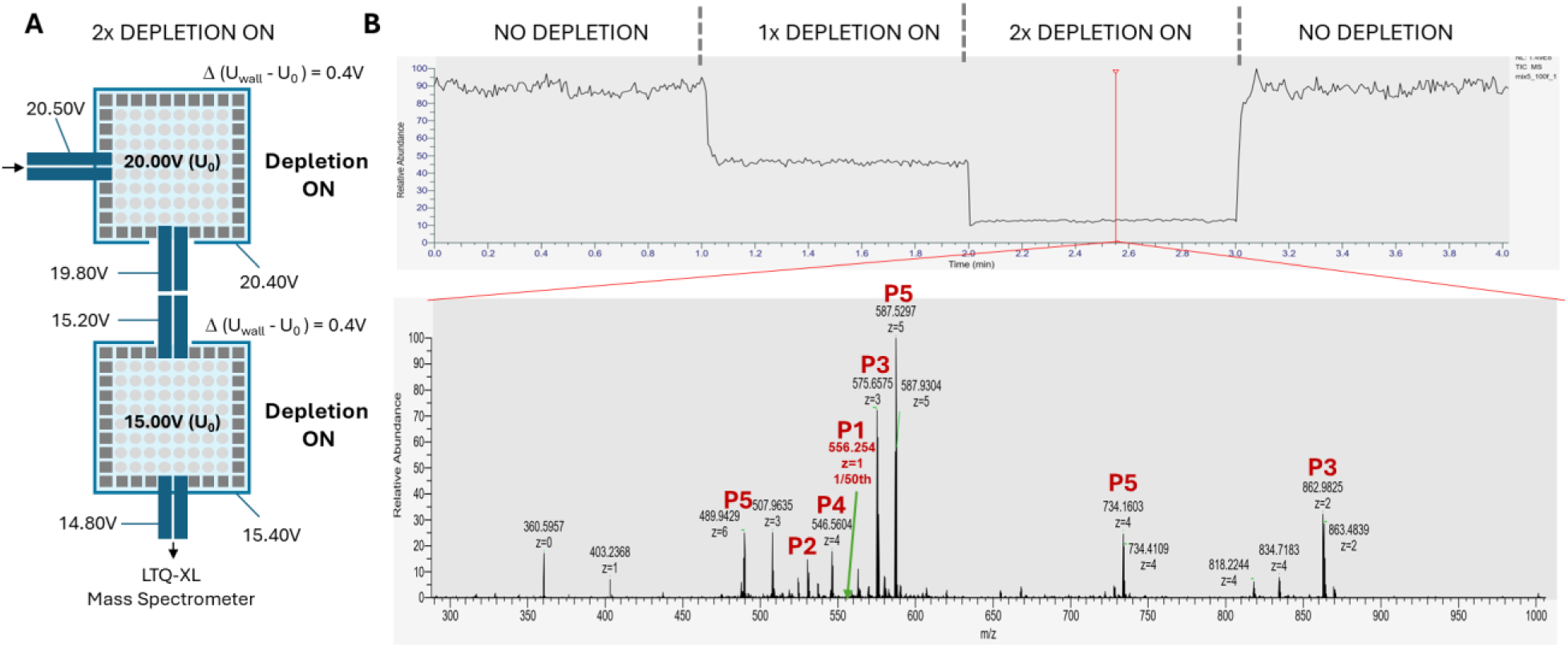
Selective depletion of singly charged ions using both traps of the tandem MultiQ-IT/LTQ-XL mass spectrometer. An equimolar mixture of 5 peptides was electrosprayed at a flow rate of 1.7 fmol/s. (**A**) Distribution of potentials in the tandem 486-quadrupole MultiQ-ITs in 2x DEPLETION ON mode, where the difference between the DC offset of both trap quadrupoles and the wall is reduced to 0.4 V. (**B**) (Top) Total ion count (TIC) and (bottom) a single-scan m/z spectrum in 2x DEPLETION ON mode. All major peaks of the 5-peptide mixture are labeled in the spectrum. The intensity of the singly charged peptide signal at m/z = 556.282 (labeled as P1) is reduced by a factor of 50 in this mode. The depletion factor can be further increased to several hundred by setting Δ(U_wall_-U_0_) to 0.3 V or smaller, albeit with some loss in total ion signal.

**Fig. S9-Part I.**
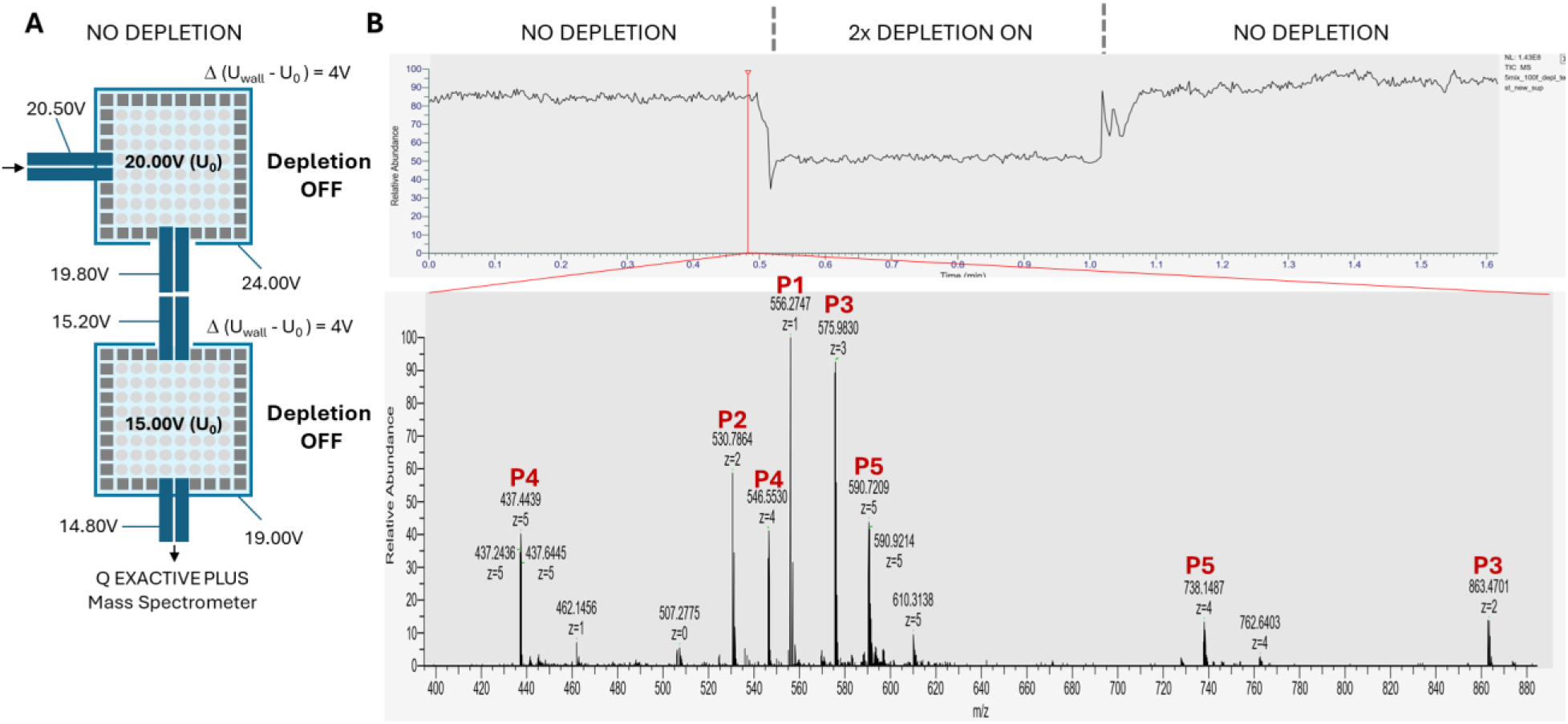
Demonstration of selective depletion of singly charged ions using the tandem MultiQ-IT/Q Exactive Plus mass spectrometer - control spectrum acquired with selective Depletion OFF is shown for comparison. An equimolar mixture of 5 peptides was electrosprayed at a flow rate of 1.7 fmol/s. (**A**) Distribution of potentials in the tandem 486-quadrupole MultiQ-ITs in NO DEPLETION mode. (**B**) (Top) Total ion count (TIC) and (bottom) a single-scan m/z spectrum in NO DEPLETION mode. All major peaks of the 5-peptide mixture are labeled in the spectrum. Note that the intensity of the singly charged peptide signal at m/z = 556.282 (labeled P1) is 100 when the tandem MultiQ-IT is in NO DEPLETION mode – i.e., Δ(U_wall_-U_o_)=4V.

**Fig. S9-Part II.**
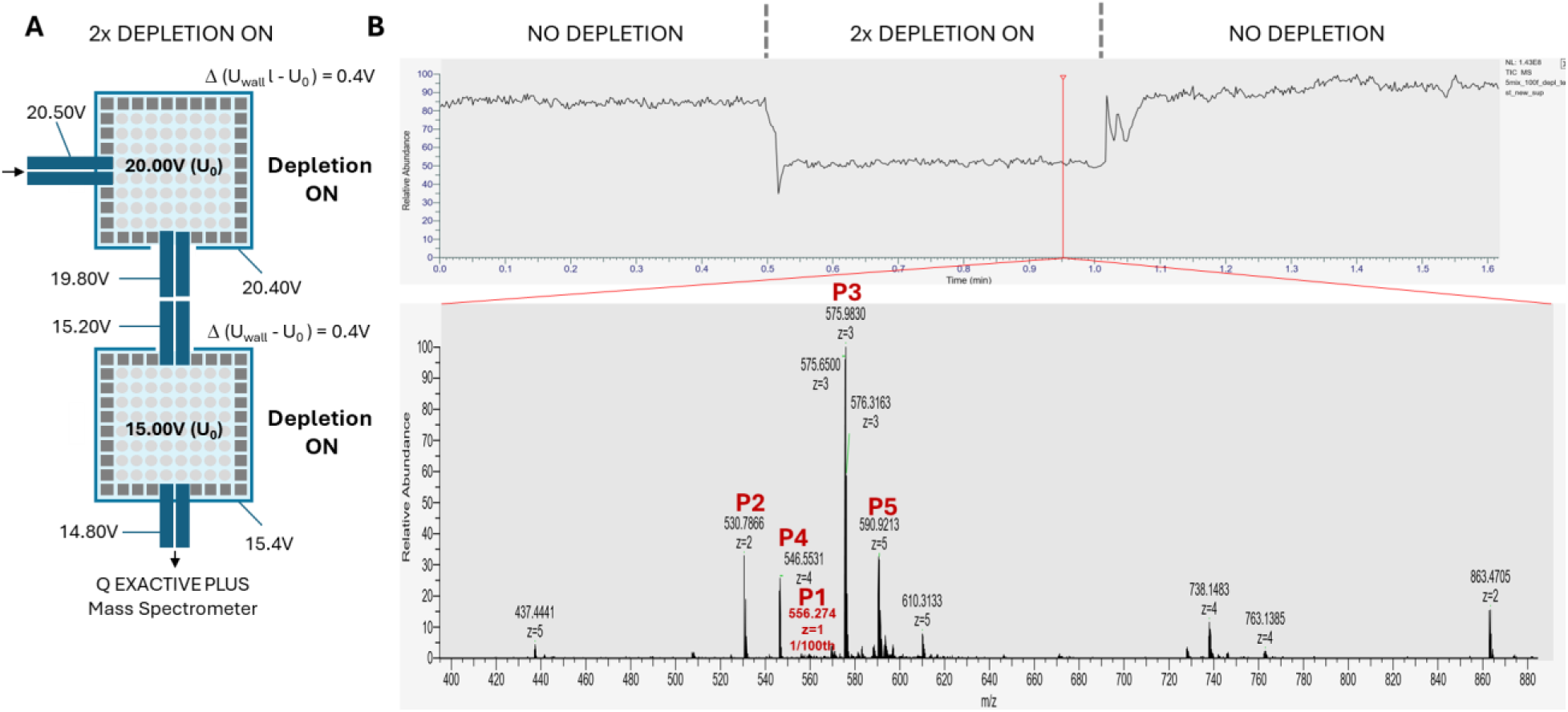
Selective depletion of singly charged ions using the tandem MultiQ-IT/Q Exactive Plus mass spectrometer. An equimolar mixture of 5 peptides was electrosprayed at a flow rate of 1.7 fmol/s. (**A**) Distribution of potentials in the tandem 486-quadrupole MultiQ-ITs in 2x DEPLETION ON mode, where the difference between the DC offset of both trap quadrupoles and the wall is reduced to 0.4 V. (**B**) (Top) Total ion count (TIC) and (bottom) a single-scan m/z spectrum in 2x DEPLETION ON mode. All major peaks of the 5-peptide mixture are labeled in the spectrum. The intensity of the singly charged peptide signal at m/z = 556.282 (labeled P1) is reduced by a factor of 100 in this mode.

**Fig. S10-Part I.**
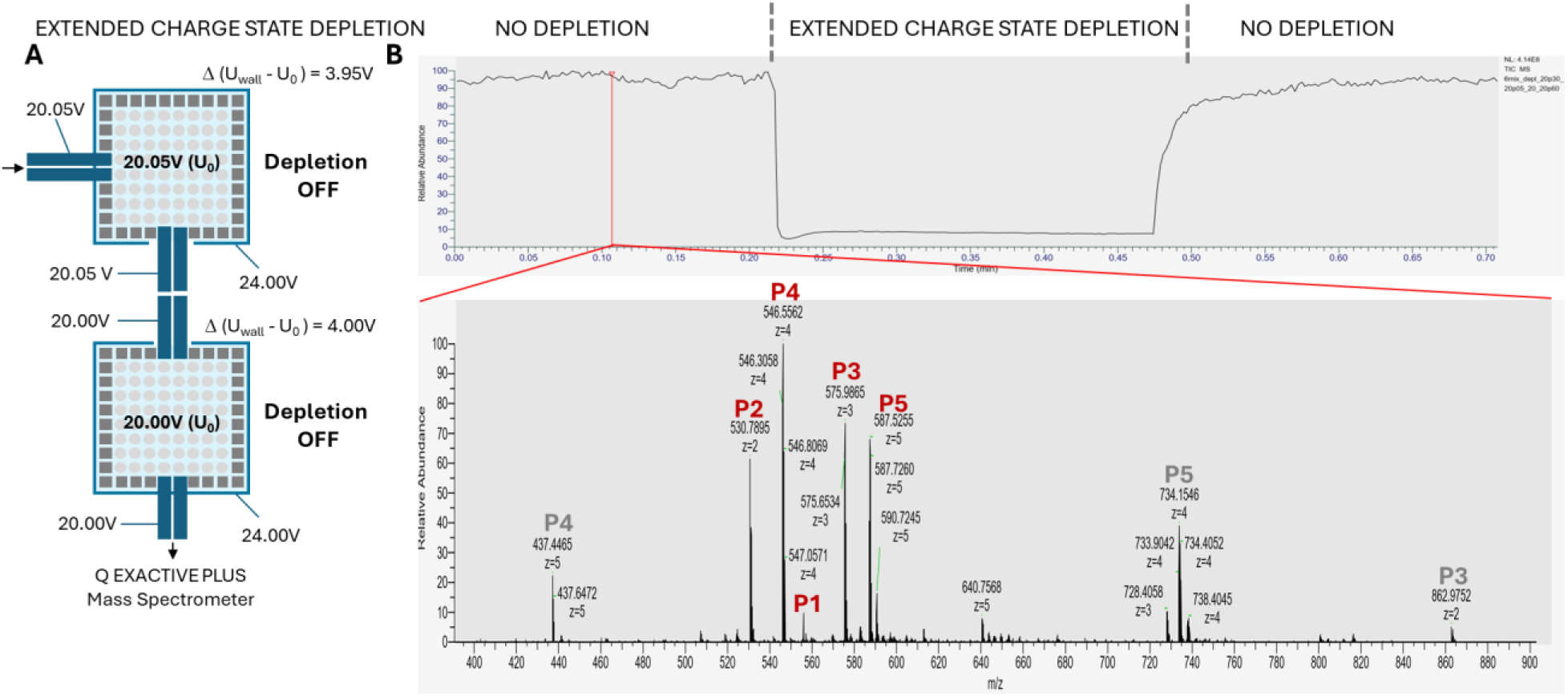
Depletion OFF control for the EXTENDED CHARGE STATE DEPLETION mode (for depleting ions in charge states z = +1, +2, and +3) in the tandem MultiQ-IT/Q Exactive Plus mass spectrometer. (**A**) Distribution of potentials in the tandem 486-quadrupole MultiQ-ITs in NO DEPLETION mode. A slightly higher offset voltage (20.05 V vs. 20.00 V) is applied to the quadrupoles of the first MultiQ-IT to create a gentle gradient for ion motion from the first to the second trap. (**B**) A total ion count (TIC) and a single-scan m/z spectrum in NO DEPLETION mode. The group of five peaks labeled at 560 ± 40 m/z labeled in red corresponds to different peptides in an equimolar mixture of 5 peptides electrosprayed at a flow rate of 1.7 fmol/s. Notably, the singly charged peptide signal at m/z = 556.277 (labeled P1) is already reduced in this mode, though its intensity should be comparable to the others (compare with fig. S9-Part I). This reduction in intensity likely originates from selective loss to the wall due to excess space charge.

**Fig. S10-Part II.**
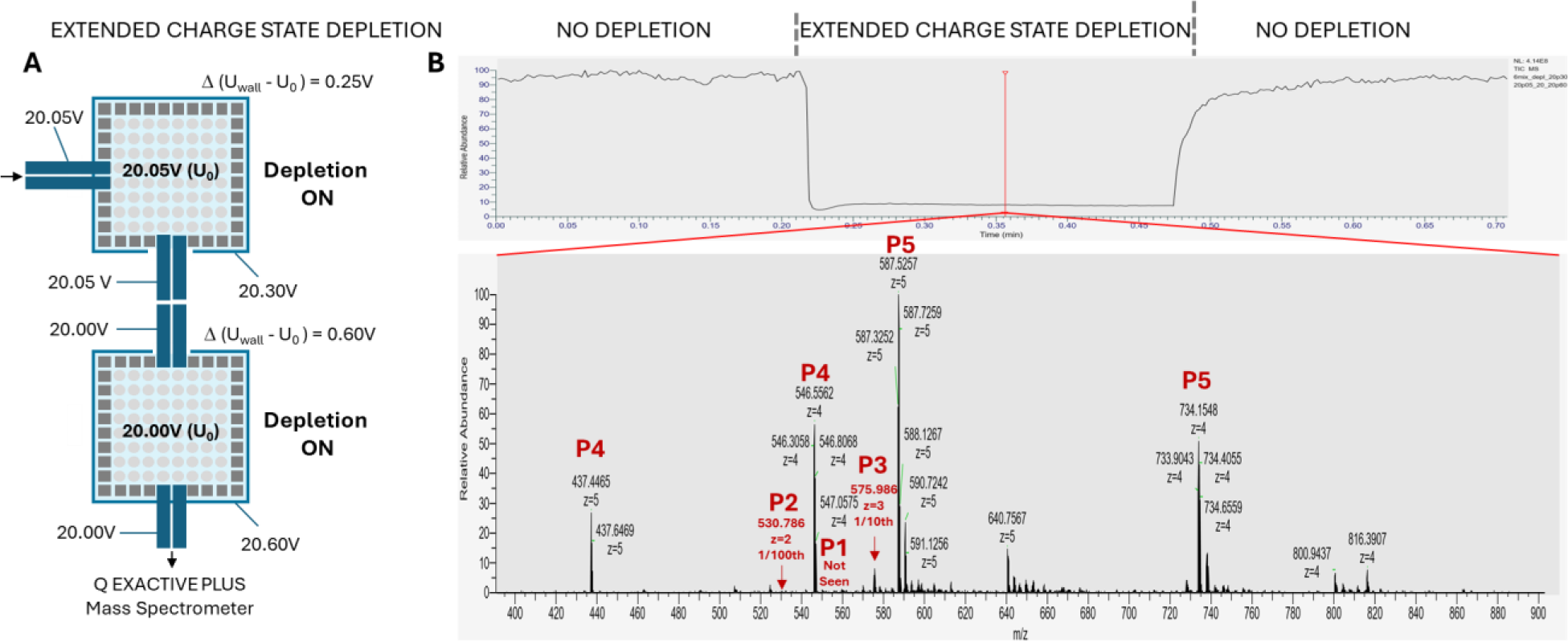
EXTENDED CHARGE STATE DEPLETION mode ON in the tandem MultiQ-IT/Q Exactive Plus mass spectrometer for depleting ions in charge states z = +1, +2, and +3. (**A**) Distribution of potentials in the EXTENDED CHARGE STATE DEPLETION mode for the tandem 486-quadrupole MultiQ-ITs. In this mode, ions pass between the two MultiQ-ITs via extended quadrupoles due to the small potential difference between the traps, enabling depletion in both. (**B**) (Top) The total ion count (TIC) and (bottom) a single-scan m/z spectrum (at the time indicated by the red line) in the EXTENDED CHARGE STATE DEPLETION mode. The group of peaks labeled at 560 ± 40 m/z labeled with red show a significant decrease in intensity for ions in charge states z = +1 (not detected), z = +2 (reduced by a factor of 100), and z = +3 (reduced by a factor of ∼10), compared to the corresponding peaks in fig. S10-Part I.

**Fig. S11-Part I.**
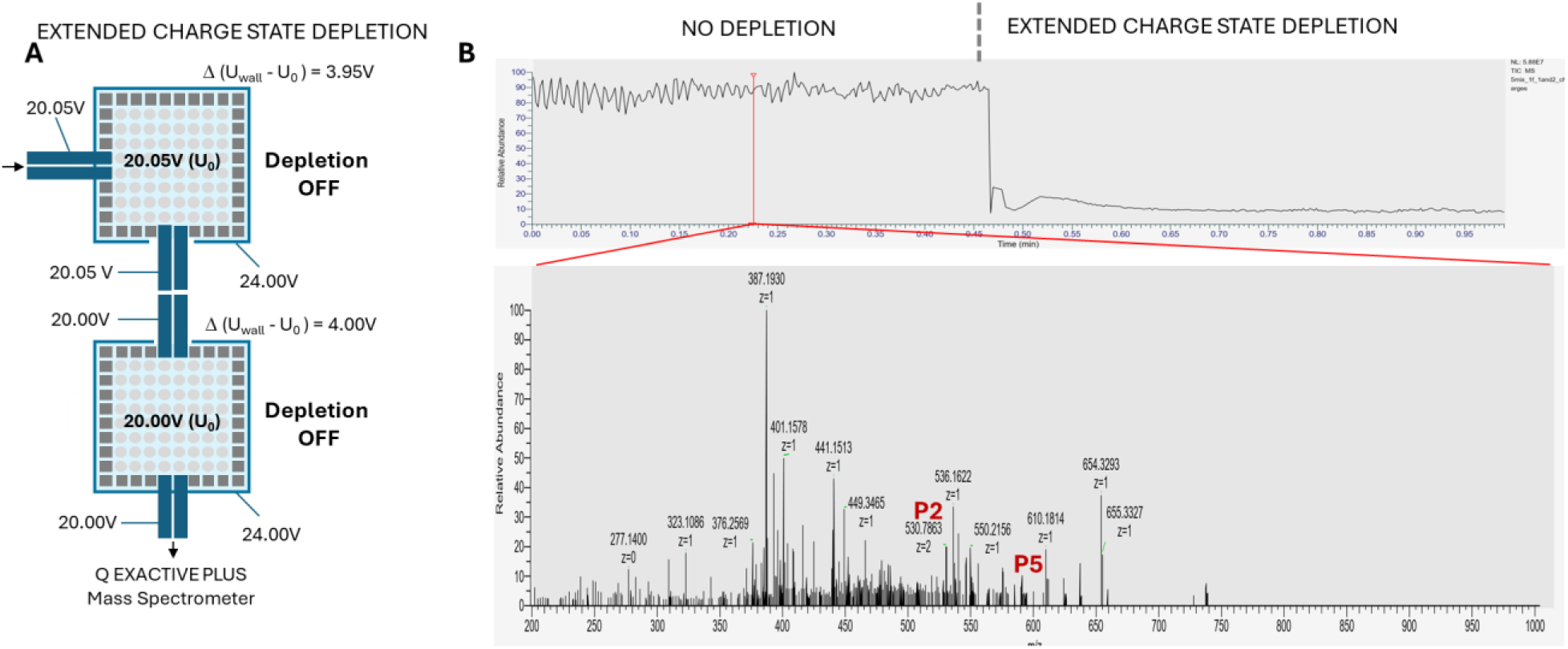
Depletion OFF control for the EXTENDED CHARGE STATE DEPLETION mode (for selectively enhancing detection of low-abundance peptide peaks with charge states z ≥ +3) in the tandem MultiQ-IT/Q Exactive Plus mass spectrometer. (**A**) Distribution of potentials in the NO DEPLETION mode for the tandem 486-quadrupole MultiQ-ITs. Ions pass between the two MultiQ-ITs via extended quadrupoles, driven by a small potential difference between traps. A slightly higher offset voltage (20.05 V vs. 20.00 V) applied to the quadrupoles of the first MultiQ-IT creates a gentle gradient for ion motion. (**B**) (Top) Total ion count (TIC) and (bottom) a single-scan m/z spectrum in NO DEPLETION mode were generated from an equimolar mixture of 5 peptides electrosprayed at a flow rate of 17 amol/s. At this low concentration, large peaks from chemical noise (see main text), obscure low-abundance peaks in the sample. Here, only two peaks from the 5-peptide mixture (see fig. S8-Part I) could barely be discerned. The next figure highlights a significant improvement in this process by using the EXTENDED CHARGE STATE DEPLETION mode.

**Fig. S11-Part II.**
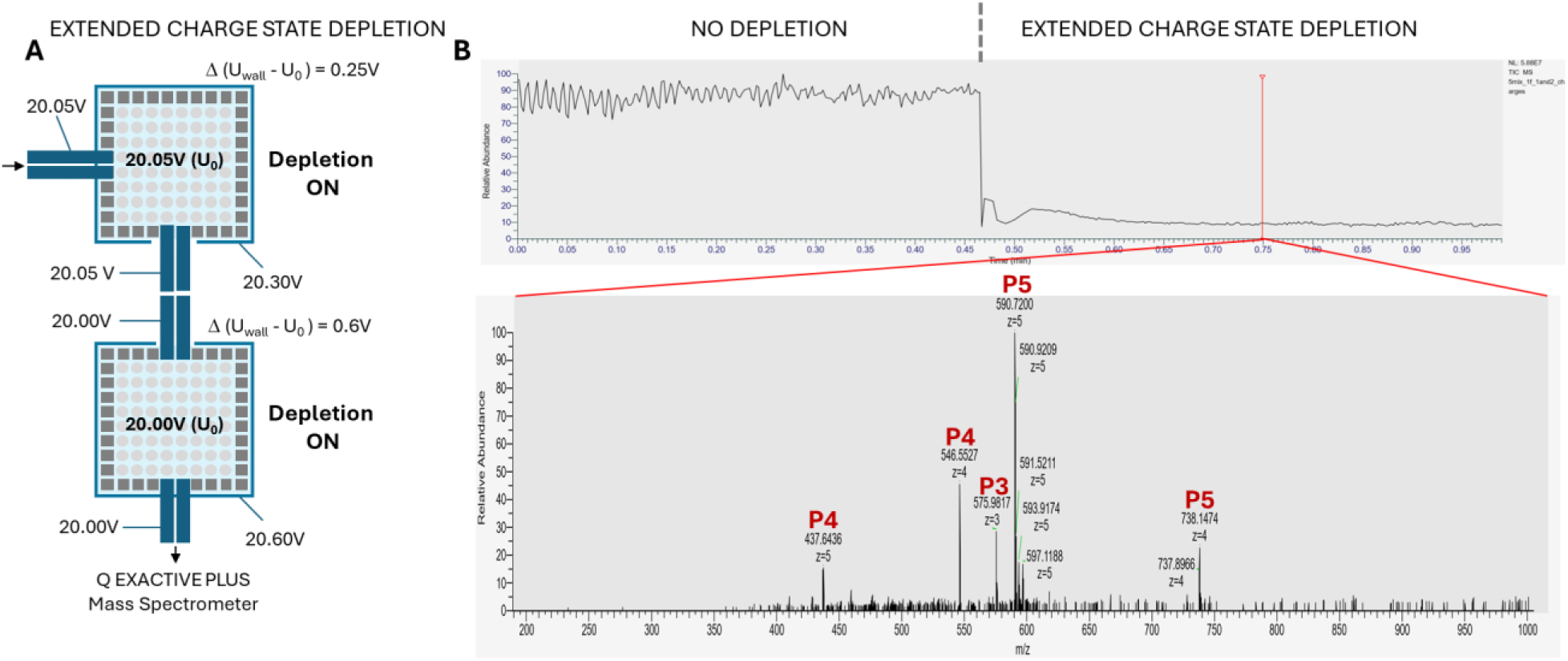
EXTENDED CHARGE STATE DEPLETION mode in the tandem MultiQ-IT/Q Exactive Plus mass spectrometer to enhance detection of low-abundance peptides with charge states z ≥ +3. (**A**) Distribution of potentials in the EXTENDED CHARGE STATE DEPLETION mode for the tandem 486-quadrupole MultiQ-ITs. Ions pass between the two MultiQ-ITs via extended quadrupoles due to a small potential difference between traps, enabling depletion in both. A slightly higher offset voltage (20.05 V vs. 20.00 V) applied to the first MultiQ-IT quadrupoles creates a gentle gradient for ion motion. (**B**) Total ion count (TIC) and a single-scan m/z spectrum in the EXTENDED CHARGE STATE DEPLETION mode were obtained from an equimolar mixture of 5 peptides electrosprayed at a flow rate of 17 amol/s. All peptide peaks with z > +3 are clearly observed, with a signal-to-noise improvement exceeding 100-fold. The remaining low-intensity background in the spectrum arises primarily from residual chemical noise with higher charged states, as demonstrated in the next two figures.

**Fig. S12-Part I.**
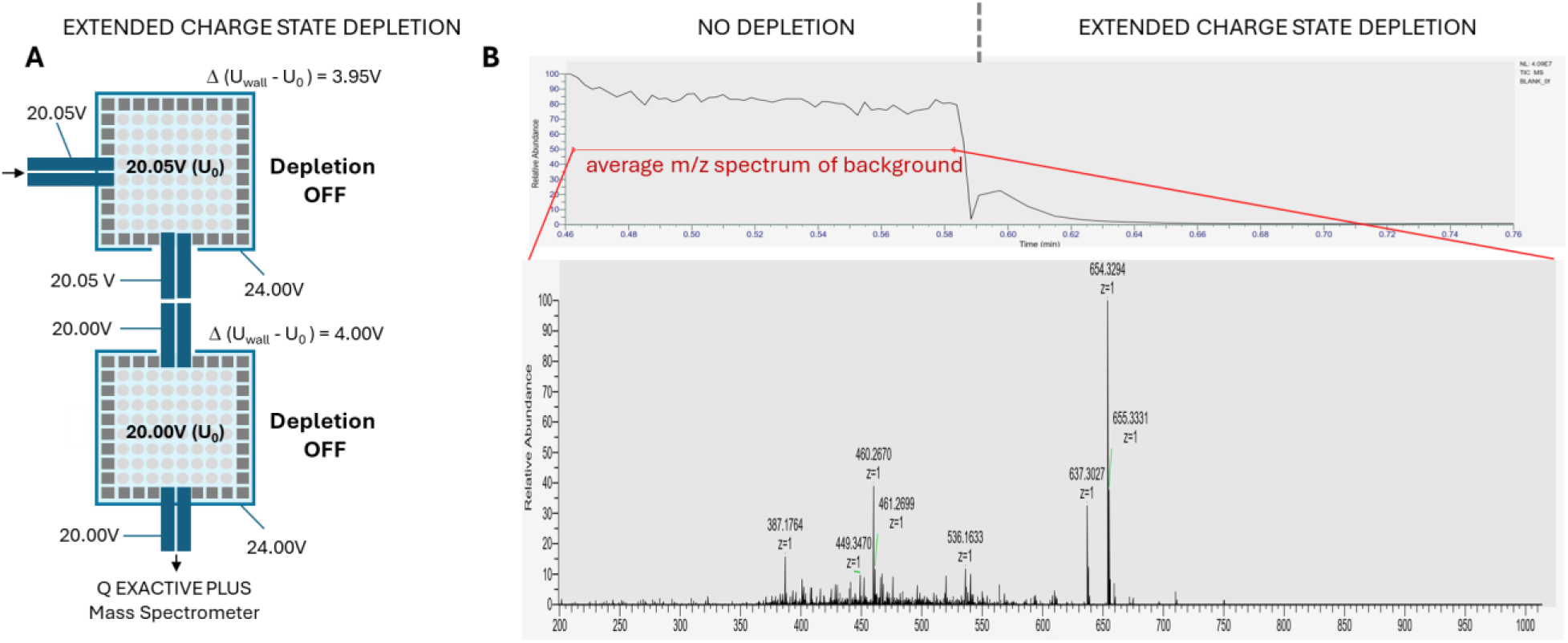
Depletion OFF control for the EXTENDED CHARGE STATE DEPLETION mode in the tandem MultiQ-IT/Q Exactive Plus mass spectrometer to investigate highly charged ions originating from chemical noise. (**A**) Distribution of potentials in NO DEPLETION mode for the tandem 486-quadrupole MultiQ-ITs. (**B**) (Top) Total ion current (TIC) and (bottom) average m/z spectrum of background in NO DEPLETION mode obtained from an acetonitrile/water/formic acid solution (30/69.9/0.1) in the absence of added sample. The prominent peaks of singly charged ions in this spectrum correspond to chemical noise (see main text). These peaks are commonly observed in the spectra of low-concentration samples.

**Fig. S12-Part II.**
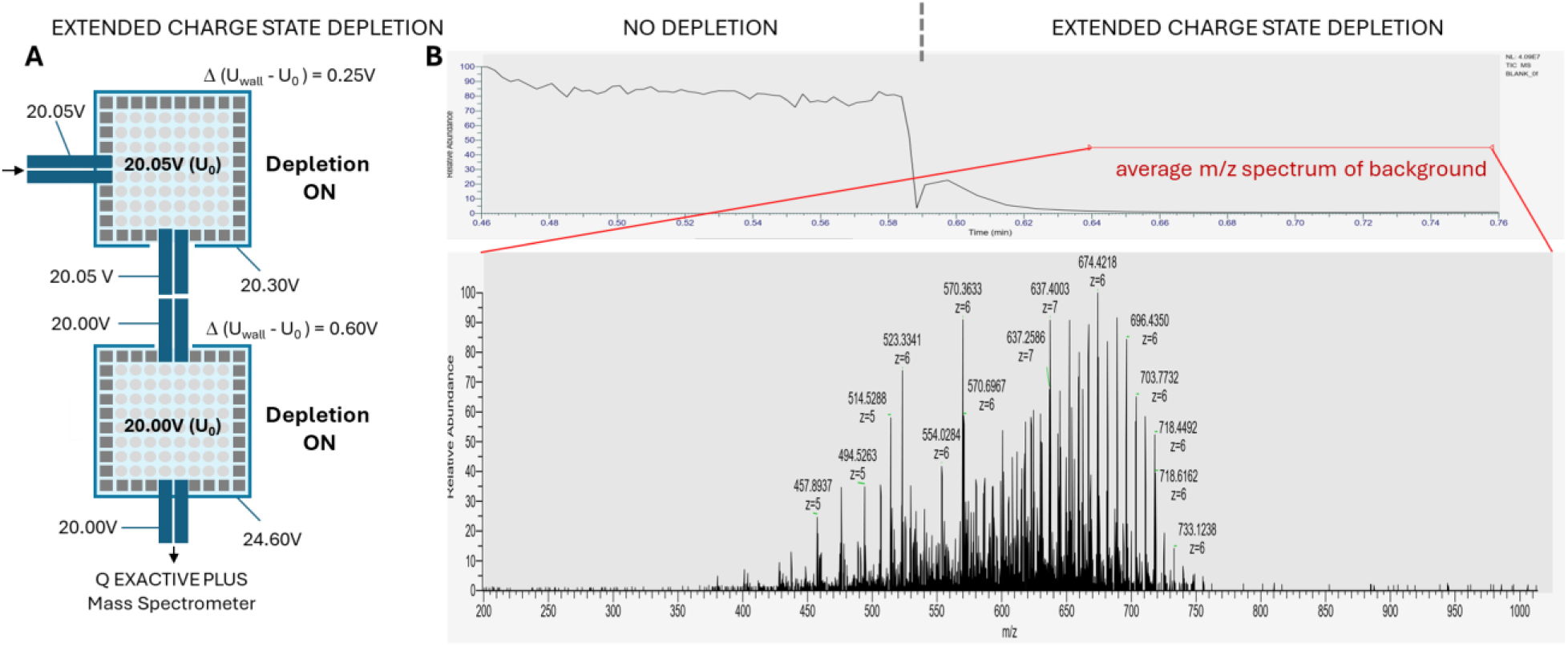
EXTENDED CHARGE STATE DEPLETION mode in the tandem MultiQ-IT/Q Exactive Plus mass spectrometer (Fig.1e) for investigating highly charged ions originating from chemical noise. No sample was added to the electrosprayed acetonitrile/water/formic acid solution (30/69.9/0.1). (**A**) In the EXTENDED CHARGE STATE DEPLETION mode, ions pass between the two MultiQ-ITs via extended quadrupoles due to a small potential difference between the traps. The higher offset voltage (20.05 V vs. 20.00 V) applied to the first MultiQ-IT quadrupoles creates a gentle gradient for ion motion. (**B**) (Top) Total ion current (TIC) and (bottom) average m/z spectra of background measured in this mode show effective depletion of z = +1, z = +2, and partial depletion of z = +3 ions, allowing explicit detection of highly charged ions. We attribute these ions to an unknown poly(ethylene glycol) (PEG)-containing compound (average molecular mass ∼4000) with multiple basic groups. The contamination was likely introduced from the polypropylene vials used for sample handling, presumably from proprietary surface treatments designed to minimize adsorption or improve surface properties. These findings underscore the importance of minimizing background contamination when analyzing small sample quantities.

**Fig. S13.**
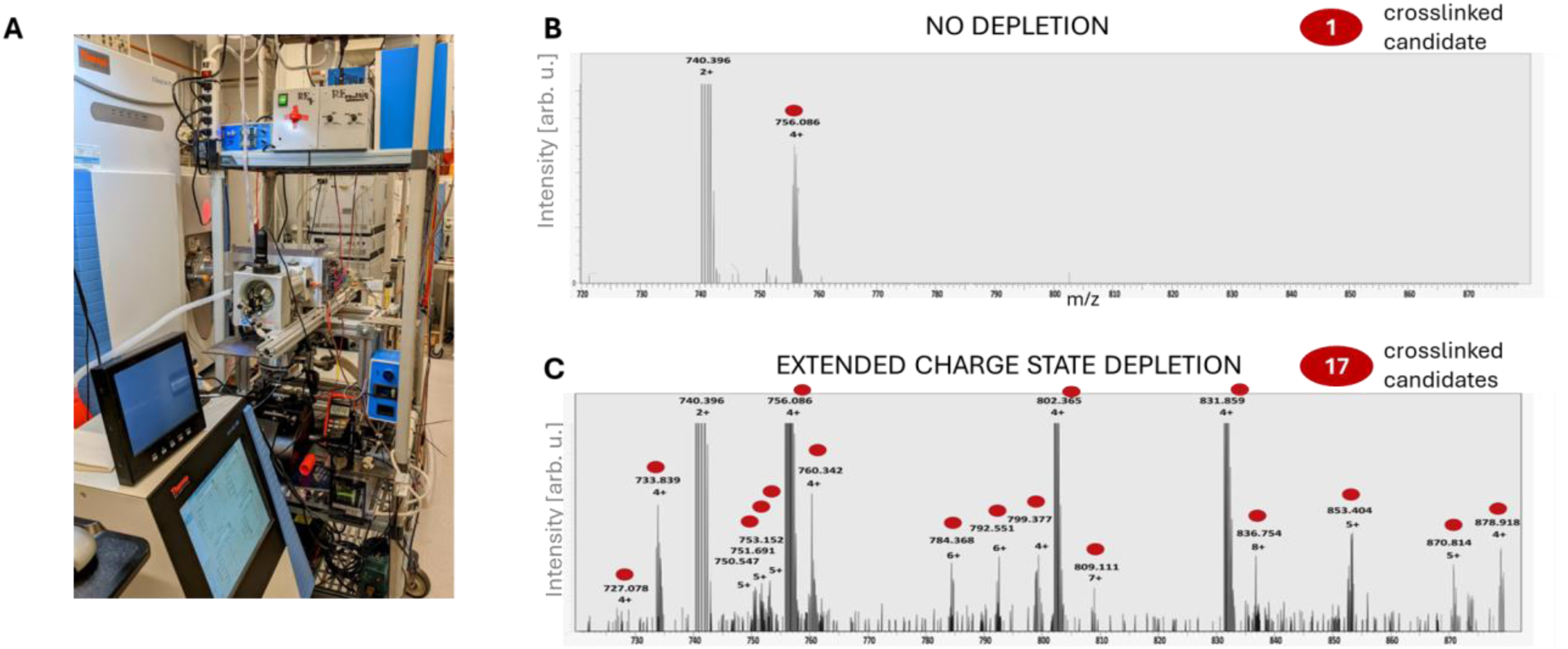
Enhancing detection of crosslinked peptide candidates by selectively depleting uncrosslinked peptides with z < +4. Tryptic digests of chemically crosslinked proteins are dominated by non-crosslinked peptides, which obscure the much lower-abundance crosslinked species. Because crosslinking is inefficient, detecting crosslinked peptides requires a mass spectrometric readout with very high dynamic range. However, the intense background of uncrosslinked peptides makes low-intensity crosslinked species difficult to detect. Since crosslinked peptides typically have charge states z ≥ +4, while uncrosslinked peptides predominantly have charge states z < +4, selective depletion of lower charge states (as demonstrated in Extended Data Figs.10–12) enriches for the crosslinked species, improving their detection. (**A**) Bovine serum albumin was crosslinked with DSS, reduced, alkylated, and digested with trypsin following standard protocols. The resulting digest was separated by HPLC (Easy-nLC 1200, Thermo) and analyzed by a tandem MultiQ-IT/Q Exactive Plus mass spectrometer. Data-dependent acquisition was used, starting with a full m/z scan followed by fragmentation of the 10 most abundant species with charge states ≥ +4, thereby excluding most non-crosslinked peptides (typically z= 2+ and 3+). (B) Portion of a mass spectrum from a single scan obtained in NO DEPLETION mode. Only a single crosslinked peptide candidate was detected, as the scan was dominated by an abundant doubly charged species (m/z = 740.396). (**C**) In contrast, the corresponding portion of the mass spectrum from the same scan obtained in EXTENDED CHARGE STATE DEPLETION mode enabled detection of 17 crosslinked peptide candidates by selectively depleting low-charge-state ions and enriching for species with z ≥ +4.

## References

1. Y. Y. Zhang, B. R. Fonslow, B. Shan, M. C. Baek, J. R. Yates, Protein Analysis by Shotgun/Bottom-up Proteomics. Chemical Reviews 113, 2343–2394 (2013).

2. R. Aebersold, M. Mann, Mass-spectrometric exploration of proteome structure and function. Nature 537, 347–355 (2016).

3. H. I. Stewart et al., Parallelized Acquisition of Orbitrap and Astral Analyzers Enables High-Throughput Quantitative Analysis. Analytical Chemistry 95, 15656–15664 (2023).

4. A. D. Brunner et al., Ultra-high sensitivity mass spectrometry quantifies single-cell proteome changes upon perturbation. Molecular Systems Biology 18, e10798 (2022).

5. R. G. Huffman et al., Prioritized mass spectrometry increases the depth, sensitivity and data completeness of single-cell proteomics. Nature Methods 20, 714–722 (2023).

6. J. L. Gustafson, Reevaluating Amdahl Law. Communications of the ACM 31, 532–533 (1988).

7. S. G. Park, G. A. Anderson, J. E. Bruce, Parallel Spectral Acquisition with Orthogonal ICR Cells. Journal of the American Society for Mass Spectrometry 28, 515–524 (2017).

8. A. A. Makarov, S. Horning, S. R. Horning, Parallel Mass Analysis. US7985950-B2, (2011).

9. A. N. Krutchinsky, V. Sherman, H. Cohen, B. T. Chait, Multi-Pole Ion Trap for Mass Spectrometry-1. US8637817-B1, (2014).

10. A. N. Krutchinsky, V. Sherman, H. Cohen, B. T. Chait, Multi-Pole Ion Trap for Mass Spectrometry-2. US8866076-B2, (2014).

11. A. N. Krutchinsky, V. Sherman, H. Cohen, B. T. Chait, Multi-Pole Ion Trap for Mass Spectrometry-3. US9129789-B2, (2015).

12. A. N. Krutchinsky, V. Sherman, H. Cohen, B. T. Chait, Multi-Pole Ion Trap for Mass Spectrometry-4. US9299550-B2, (2016).

13. M. P. Rout et al., The yeast nuclear pore complex: Composition, architecture, and transport mechanism. Journal of Cell Biology 148, 635–651 (2000).

14. M. P. Rout, J. D. Aitchison, M. O. Magnasco, B. T. Chait, Virtual gating and nuclear transport: the hole picture. Trends in Cell Biology 13, 622–628 (2003).

15. J. Franzen, M. Schubert, Method and device for the reflection of charged particles on surfaces. US5572035-A, (1996).

16. C. M. Whitehouse, D. G. Welkie, L. Cousins, RF surfaces and RF ion guides. US7365317-B2, (2008).

17. R. E. March, J. F. J. Todd, Practical Aspects of Ion Trap Mass Spectrometry Volume 1: Fundamentals of Ion Trap Mass Spectrometry. T. Cairns, Ed., CRC Series Modern Mass Spectrometry (CRC Press, Boca Raton New York London Tokyo, 1995).

18. A. E. Holme, Operation of a quadrupole mass filter with only an RF voltage component applied to rod system. International Journal of Mass Spectrometry and Ion Processes 22, 1–5 (1976).

19. U. Brinkmann, A modified quadrupole mass filter for the separation of ions of higher masses with high transmission. International Journal of Mass Spectrometry and Ion Physics 9, 161–166 (1972).

20. F. A. Londry, J. W. Hager, Mass selective axial ion ejection from a linear quadrupole ion trap. Journal of the American Society for Mass Spectrometry 14, 1130–1147 (2003).

21. S. K. Chowdhury, V. Katta, B. T. Chait, An electrospray ionization mass spectrometer with new features. Rapid Communications in Mass Spectrometry 4, 81–87 (1990).

22. S. K. Chowdhury, V. Katta, B. T. Chait, Electrospray ionization mass spectrometer with new features. US4977320, (1990).

23. C. M. Rose et al., Multipurpose Dissociation Cell for Enhanced ETD of Intact Protein Species. Journal of the American Society for Mass Spectrometry 24, 816–827 (2013).

24. N. M. Riley et al., Enhanced Dissociation of Intact Proteins with High Capacity Electron Transfer Dissociation. Journal of the American Society for Mass Spectrometry 27, 520–531 (2016).

25. R. A. Zubarev, A. Makarov, Orbitrap Mass Spectrometry. Analytical Chemistry 85, 5288–5296 (2013).

26. D. Gerlich, Inhomogenous RF-fields - A versatile tool for the study of processes with slow ions. Advances in Chemical Physics 82, 1–176 (1992).

27. J. W. Hager, Method of mass spectrometry, to enhance seperation of ions with different charges. US2004183005-A1, (2004).

28. A. V. Tolmachev, A. N. Vilkov, L. Pasa-Tolic, H. R. Udseth, R. D. Smith, Suppression of the lower charge state ions in the external accumulation RF multipole with a reduced trapping DC potential. Journal of the American Society for Mass Spectrometry 14, 1229–1235 (2003).

29. H. C. Berg, Random Walks in Biology, Expanded edition. (Princeton University Press, 1983).

30. A. N. Krutchinsky, B. T. Chait, On the nature of the chemical noise in MALDI mass spectra. Journal of the American Society for Mass Spectrometry 13, 129–134 (2002).

31. A. Maiolica et al., Structural analysis of multiprotein complexes by cross-linking, mass spectrometry, and database searching. Molecular & Cellular Proteomics 6, 2200–2211 (2007).

32. A. Leitner et al., Probing Native Protein Structures by Chemical Cross-linking, Mass Spectrometry, and Bioinformatics. Molecular & Cellular Proteomics 9, 1634–1649 (2010).

33. C. W. Akey et al., Comprehensive structure and functional adaptations of the yeast nuclear pore complex. Cell 185, 361–378 (2022).

34. F. Alber et al., The molecular architecture of the nuclear pore complex. Nature 450, 695–701 (2007).

35. F. Alber et al., Determining the architectures of macromolecular assemblies. Nature 450, 683–694 (2007).

36. S. J. Kim et al., Integrative structure and functional anatomy of a nuclear pore complex. Nature 555, 475–482 (2018).

37. X. H. Zhang et al., Reducing Space Charge Effects in a Linear Ion Trap by Rhombic Ion Excitation and Ejection. Journal of the American Society for Mass Spectrometry 27, 1256–1262 (2016).

38. M. Winey, D. Yarar, T. H. Giddings, D. N. Mastronarde, Nuclear pore complex number and distribution throughout the Saccharomyces cerevisiae cell cycle by three-dimensional reconstruction from electron micrographs of nuclear envelopes. Molecular Biology of the Cell 8, 2119–2132 (1997).

39. P. Mielczarek, J. Silberring, M. Smoluch, Minituarization in mass spectrometry. Mass Spectrometry Reviews 39, 453–470 (2020).

